# Pharmacological control of CAR T cells through CRISPR-driven rapamycin resistance

**DOI:** 10.1101/2023.09.14.557485

**Authors:** Sébastien Levesque, Gillian Carleton, Victoria Duque, Claudia Goupil, Jean-Philippe Fiset, Sarah Villeneuve, Eric Normandeau, Geneviève Morin, Nellie Dumont, Brad H. Nelson, Josée Laganière, Brian Boyle, Julian J. Lum, Yannick Doyon

## Abstract

Chimeric antigen receptors (CARs) reprogram T cells to recognize and target cancer cells. Despite remarkable responses observed with CAR-T cell therapy in patients with hematological malignancies, CAR-T cell engineering still relies mostly on randomly integrating vectors, limiting the possibilities of fine-tuning T cell function. Here, we designed a CRISPR-based marker-free selection strategy to simultaneously target a therapeutic transgene and a gain-of-function mutation to the *MTOR* locus to enrich cells resistant to rapamycin, a clinically used immunosuppressant. We readily engineered rapamycin-resistant (RapaR) CAR-T cells by targeting CAR expression cassettes to the *MTOR* locus. Using *in vitro* cytotoxicity assays, and a humanized mouse model of acute lymphoblastic leukemia, we show that RapaR-CAR-T cells can efficiently target CD19^+^ leukemia cells in presence of immunosuppressing doses of rapamycin. Furthermore, our strategy allows multiplexed targeting of rapamycin-regulated immunoreceptors complexes (DARICs) to the *MTOR* and *TRAC* loci to pharmacologically control CAR-T cells’ activity. We foresee that our approach could both facilitate the enrichment of CRISPR-engineered CAR-T cells *ex vivo* and *in vivo* while improving tumor eradication.

## INTRODUCTION

Adoptive cell therapy with chimeric antigen receptor T (CAR-T) cells has shown remarkable potency against relapsed and refractory acute lymphoblastic leukemia (ALL) and non-Hodgkin lymphoma^1–5^. CAR-T cells are genetically engineered T cells that are able to recognize and target cancer cells expressing a specific antigen on their surface^6, 7^. Randomly integrating vectors based on retroviruses, lentiviruses or *piggyBac* transposons are routinely used for cell therapy applications, but genome editing technologies offer a wide a range of opportunities to enhance CAR-T cells functionality^8, 9^. The promising potential of CRISPR-based technologies to engineer CAR-T cells has been demonstrated by targeting a CAR to the *TRAC* locus to avert tonic CAR signaling and delay T cells differentiation and exhaustion^10^, or by generating HLA-independent T cell receptors (HIT) to afford high antigen sensitivity^11^. CRISPR-based technologies also offer great potential to inactivate key genes involved in T cells dysfunction^12–14^ or to generate allogeneic universal CAR-T cells^15–17^.

While progress has been made towards high-yield targeted transgene integration in primary cells^10, 11, 18^, enrichment strategies could further facilitate the development of CRISPR-engineered cell therapies. To enrich targeted transgene integration via homology-directed repair (HDR) in primary T cells, a selection method based on the expression of a methotrexate-resistant dihydrofolate reductase mutant has previously been reported^19^. A metabolic safety switch has also been developed via *UMPS* knockout to enrich T cells expressing reporter transgenes^20^. However, no marker-free selection strategy has been developed to simultaneously enrich CRISPR-engineered CAR-T cells and enhance their antitumor activity.

The serine/threonine protein kinase mTOR constitutes the catalytic subunit of two distinct complexes, known as mTORC1 and mTORC2, which coordinate eukaryotic cell growth and metabolism by integrating a diverse set of environmental inputs^21–23^. mTORC1 controls cellular growth and metabolism by sensing nutrient levels and growth factor signals to coordinate anabolic and catabolic metabolism^24^. While mTORC1 is sensitive to rapamycin, an immunosuppressant also used in oncology, mTORC2 remains functional upon acute rapamycin treatment^21–23^. Considering that mTOR signaling is commonly activated in tumor cells and alters the expression and the activity of key metabolic enzymes^21, 25, 26^, combinatorial CAR-T cell therapy with rapamycin could synergistically enhance antitumor activity. Indeed, random integration of a rapamycin-resistant mTOR transgene and a CD19-CAR in primary T cells using *piggyBac* transposons has been shown to confer a selective growth advantage in the presence of rapamycin and enhanced antitumor activity when used in combination^27^. However, delivery of this large transgene is limiting, and product-derived lymphomas have been reported with *piggyBac*-modified CAR-T cells^28, 29^, so editing the endogenous *MTOR* locus to generate dominant cellular resistance to rapamycin using CRISPR could be a viable alternative. Since mTORC1 signaling lies downstream of several activation signals essential for T cell activation and expansion^21, 30^, this central signaling node is a prime target to develop a tissue-specific marker-free selection strategy for CRISPR-engineered CAR-T cells. More importantly, CRISPR-induced drug resistance offers great potential for pharmacological control of CAR-T cells activity through receptor dimerization^31, 32^ even when using an immunosuppressive dose of rapamycin.

In this study, we performed saturation prime editing^33^ at the *MTOR* locus to identify and characterize optimal mutations conferring rapamycin resistance. We then devised an intron nesting strategy^34, 35^ to enrich CRISPR-engineered cells *ex vivo* and *in vivo* without using an exogenous selection marker. We demonstrate that targeting CAR expression cassettes to the *MTOR* locus allows the selection and regulation of rapamycin-resistant CAR-T cells to perform combinatorial immunotherapy with rapamycin. More broadly, we propose a versatile approach to facilitate targeted therapeutic transgene integration for cell therapy applications.

## RESULTS

### Identification and characterization of mutations conferring rapamycin resistance via saturation prime editing

After an extraordinary 18-month response to everolimus, a rapamycin analog, a patient with metastatic anaplastic thyroid carcinoma relapsed with a resistant tumor harboring the *MTOR*- F2108L mutation^36^. The same mutation was also found in spontaneous rapamycin-resistant yeast mutants^37^, and a clonal MCF-7 breast cancer cell line after three months of exposure to rapamycin^38^. Of importance, this mutation does not hyperactivate mTOR^34, 36, 38^ and confers resistance to first-generation allosteric inhibitors of mTOR, but not to second- and third-generation inhibitors^36, 38^. F2108 lies within the FKBP12-rapamycin binding (FRB) domain of mTOR and is an important residue involved in rapamycin interactions^39^. We performed saturation mutagenesis of F2108 to install all possible amino acids substitutions at the endogenous *MTOR* locus to identify and characterize alternative mutations causing rapamycin-resistance at this position. We devised a saturation prime editing (PE) screen^33^ using K562 cells stably expressing the mScarlet-I mTOR signaling indicator (mSc-TOSI)^34, 40^ as a phenotypic readout to interrogate the functional impact of these mutations (**Fig. 1a,b** and **Supplementary Fig. 1**). Under active mTORC1 signaling, the fluorescent mSc-TOSI reporter is rapidly phosphorylated by S6K, ubiquitinated, and degraded by the proteasome while mTOR inhibition stabilizes the reporter (**Fig. 1b**)^34, 40^.

**Figure 1.**
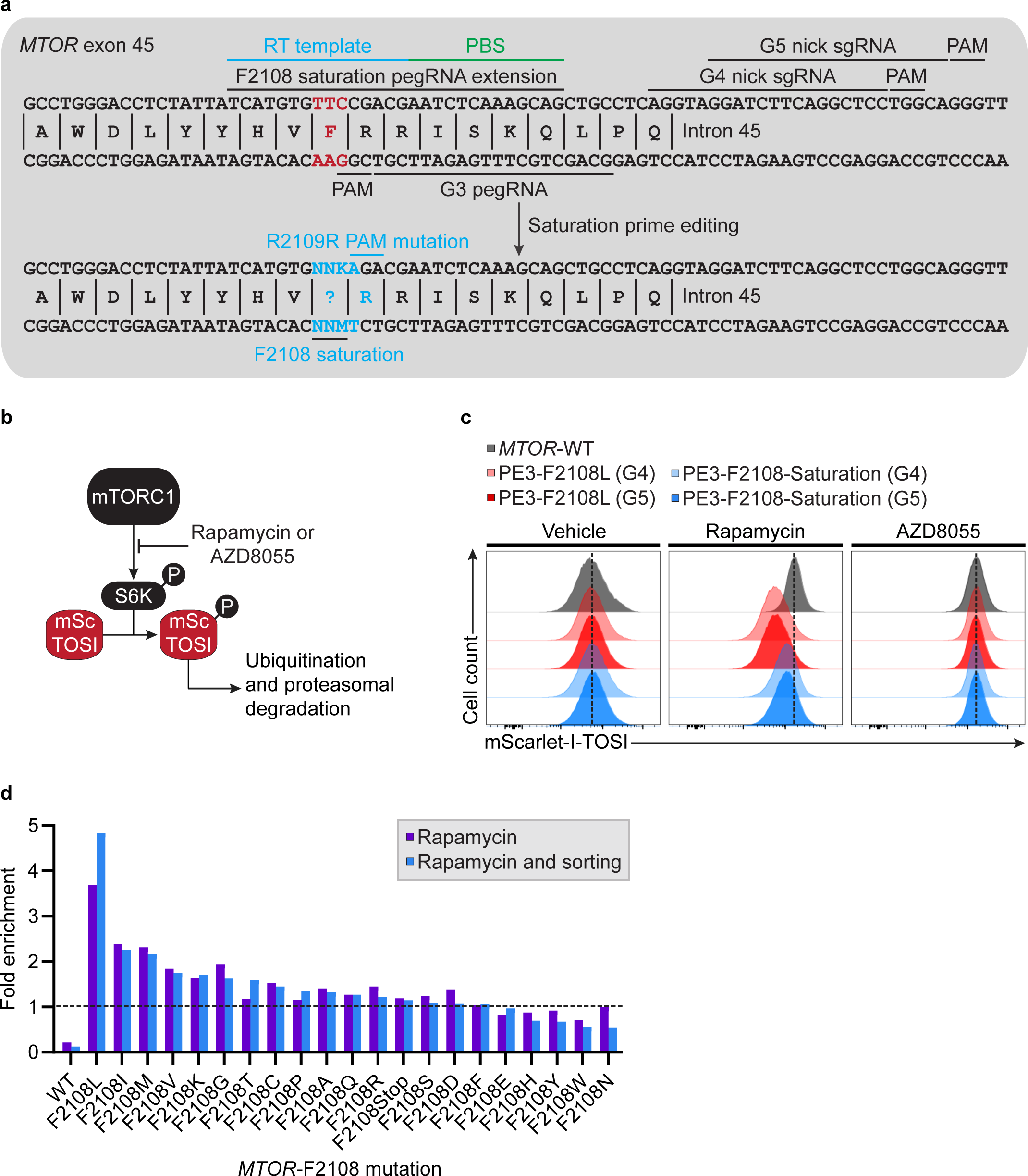
Identification of mutations conferring rapamycin resistance via saturation prime editing. (**a**) Schematic representation of the SpCas9 target sites, the reverse transcriptase (RT) template and the primer binding site (PBS) used to perform genome editing at *MTOR*. To perform saturation prime editing, G3 is used as a pegRNA, and G4 or G5 are used as a complementary nick sgRNA (PE3). The *MTOR*-F2108 codon is mutated to NNK and an additional silent PAM mutation is introduced. (**b**) Schematic of mScarlet-I mTOR signaling indicator (mSc-TOSI) degradation under mTORC1 signaling. (**c**) Histogram plot of mSc-TOSI intensity in bulk populations of cells harboring different *MTOR*-F2108 mutations. Where indicated, cells were treated for 24 hours with 50 nM rapamycin or 50 nM AZD8055 before FACS analysis. Representative images are from one of two independent biological replicates performed at different times with equivalent results. (**d**) High throughput sequencing quantification of the *MTOR*-F2108 mutations introduced via saturation prime editing following rapamycin selection and FACS sorting. The fold enrichment of each *MTOR*-F2108 mutation was calculated from the before selection sample.

We transfected our reporter cell line with a library of engineered pegRNA (epegRNA)^41^ encoding F2108 variants and performed rapamycin selection starting 3 days post-transfection until all non-resistant cells were eliminated. Selection was performed at 0.5 µM rapamycin, the lowest concentration of this drug that strongly abrogates K562 cell proliferation under our growth conditions. We performed high-throughput sequencing prior and after FACS-based cell sorting and analyzed the fold enrichment of all possible *MTOR*-F2108 permutations (**Fig. 1c,d** and **Supplementary Fig. 1**). The most enriched mutation was F2108L^36, 38^ (4.8 fold enrichment), but five additional mutations displaying 1.6 to 2.3 fold enrichments were selected for further characterization (**Fig. 1d**). Of importance, cells modified at *MTOR* remained sensitive to the 2^nd^ generation ATP-competitive inhibitor AZD8055^42^, which could be used as a safety switch in case of overt toxicity (**Fig. 1c** and **Supplementary Fig. 1** and **2**).

To validate the top hits, we designed epegRNAs to test these variants individually. K562 cells harboring these *MTOR*-F2108 substitutions grew robustly in the presence of 0.5 µM rapamycin and we observed a marked increase in the percentage of alleles harboring the PE-specified edits after rapamycin selection (**Supplementary Fig. 1**). As expected, no resistance and no enrichment were observed with the silent F2108F mutation (**Supplementary Fig. 1**). These observations confirmed that the five new variants identified by saturation prime editing conferred dominant cellular resistance to rapamycin. Furthermore, mTORC1 signaling remained fully functional with the F2108L mutation but was completely blocked with the silent F2108F mutation (**Supplementary Fig. 1 and 2**). However, intermediate levels of signaling in the presence of rapamycin were observed with F2108I, F2108M, and F2108G while weak activity was observed with F2108V and F2108K (**Supplementary Fig. 1 and 2**). These observations suggest that robust K562 cell growth can be maintained with partial mTORC1 signaling. These *MTOR*-F2108 mutations could potentially be used to pharmacologically fine-tune mTORC1 signaling while maintaining robust cellular growth. Overall, mTORC1 signaling remains completely functional in the presence of rapamycin with F2108L, and this mutation was selected to develop a marker-free selection strategy to engineer rapamycin-resistant CAR-T cells.

### Concomitant transgene integration and induction of rapamycin resistance by nuclease-driven *MTOR* editing in primary T cells

While creating F2108L by itself via PE could be used to potentiate CAR-T cells, we aimed to determine if we could couple rapamycin resistance to transgene integration in a single step by editing the *MTOR* locus via homology-directed repair (HDR). As we previously showed for the essential *ATP1A1* gene^34^, this strategy can work by nesting a transgene within an intron without disrupting the recipient gene^34, 35^. We first designed two sgRNAs targeting *MTOR* intron 45 in proximity to F2108 using CRISPOR^43^ (see G4 and G5 in **Fig. 1a**). These two chemically modified sgRNAs generated 86-90% indels in primary T cells when delivered as RNPs with both SpCas9 and the high-fidelity variant HiFiCas9^44^ (**Fig. 2a**). To assess their genome-wide specificity, we performed GUIDE-Seq^45, 46^ in primary CD3^+^ T cells from two healthy donors using wild-type SpCas9. While 9 potential off-target sites were identified with *MTOR*-G5, only one was identified with *MTOR*-G4 in CD3^+^ T cells (**Fig. 2b** and **Supplementary Fig. 3**). Similar results were obtained in K562 cells via plasmid overexpression (**Supplementary Fig. 3**). We then performed RNPs nucleofection in CD3^+^ T cells from two healthy donors followed by amplicon sequencing to validate GUIDE-Seq off-targets and test whether HiFiCas9 could mitigate off-target activity. While the frequency of alleles with indels above the 0.1% limit of detection was observed at all off-targets with SpCas9, HiFiCas9 abrogated off-target editing at all off-targets except *MTOR*-G5 OT1 (**Fig. 2c**). Since *MTOR*-G4 is highly active and specific, we prioritized this sgRNA for our studies.

**Figure 2.**
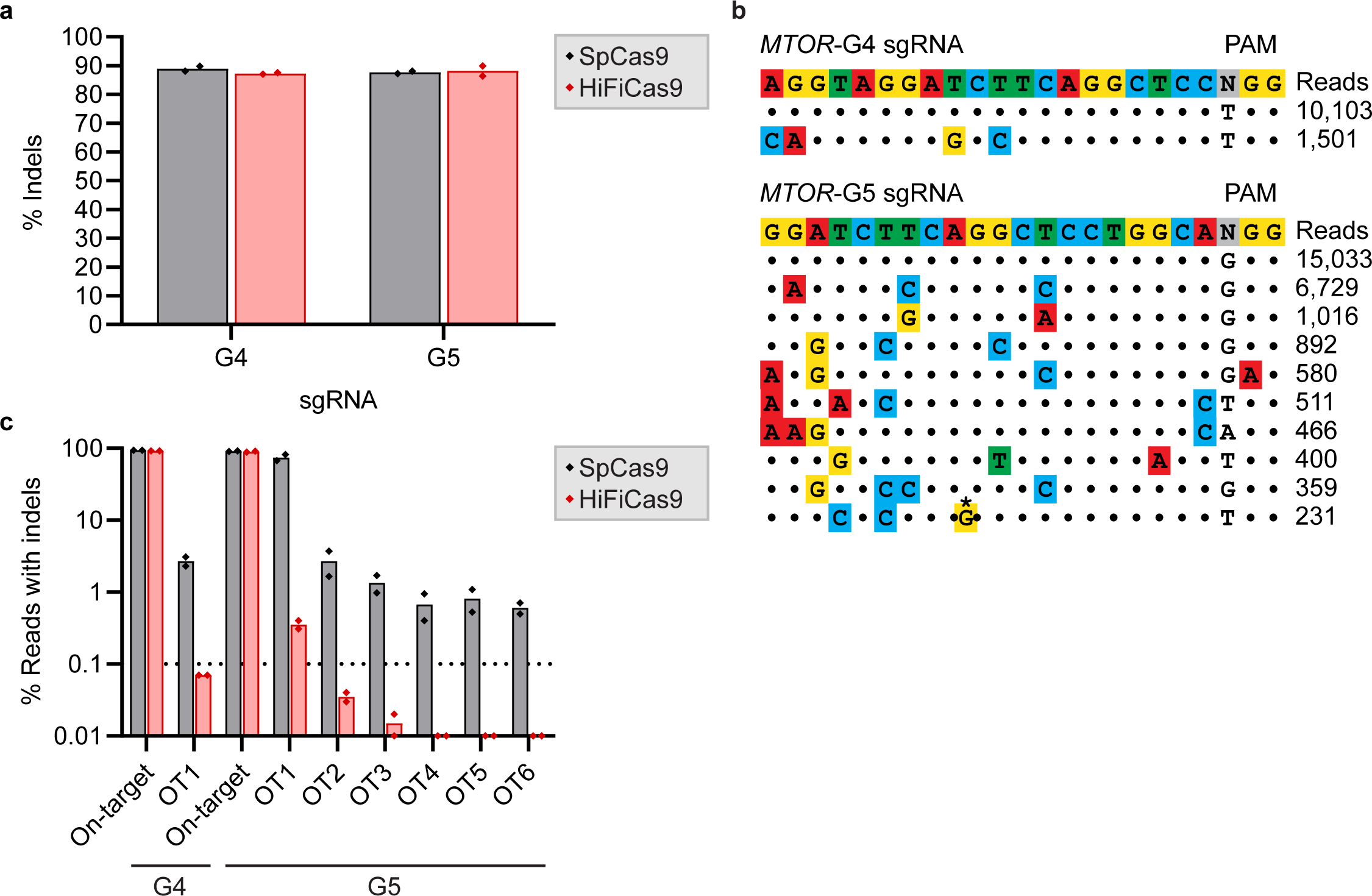
Identification of highly active and specific sgRNAs targeting the *MTOR* locus. (**a**) Small indels quantification as determined by TIDE analysis from Sanger sequencing. Primary CD3^+^ and CD8^+^ T cells were transfected with the indicated nuclease/sgRNA RNPs and genomic DNA was harvested 3 days post-transfection. *n* = 2 independent biological replicates performed at different times with CD3^+^ and CD8^+^ T cells from two different healthy donors. (**b**) Candidate off-target sites identified using GUIDE-Seq. CD3^+^ T cells were transfected with SpCas9 RNPs targeting *MTOR* intron 45 and GUIDE-Seq dsDNA. Four days post-nucleofection, CD3^+^ T cells were restimulated and expanded for an additional 7 days. Genomic DNA was harvested after expansion (11 days post-nucleofection). Dots represent matches with the intended target sequence, mismatches are colored, and nucleotide bulges are highlighted with a star. GUIDE-Seq read counts are from one of two independent biological replicates performed with CD3^+^ T cells from two different donors (see **Supplementary Fig. 3**). (**c**) Small indels quantification as determined by CRISPResso2 analysis from amplicon sequencing. Primary CD3^+^ T cells were transfected with the indicated nuclease/sgRNA RNPs and genomic DNA was harvested 3 days post-transfection. *n* = 2 independent biological replicates performed with CD3^+^ T cells from two different donors.

Next, using Cas9 RNPs and PCR-generated linear dsDNA donors^47^, we confirmed that rapamycin resistance could be achieved in primary CD3^+^ T cells and observed a marked increase to up to 62% of alleles harboring the *MTOR*-F2108L mutation after selection (**Supplementary Fig. 4**). This increase in HDR in the population upon selection was concomitant with a decrease of indels detected at the cut site, suggesting efficient elimination of cells not having undergone HDR (**Supplementary Fig. 4**). No enrichment was observed when using a donor that harbored the wildtype *MTOR*-F2108 codon, confirming that robust growth in the presence of rapamycin depends on the *MTOR*-F2108L mutation (**Supplementary Fig. 4**). In these experiments, a dose of 25 nM rapamycin was chosen based on growth curves indicating that maximal inhibition occurred in the 16-32 nM range under our conditions (data not shown). A similar dose was used in previous studies^27^ and it falls between the immunosuppressive window of 3-22 nM and the maximal steady-state plasma drug concentrations observed in clinical settings^31, 48^. Thus, the installation of the well-characterized *MTOR*-F2108L mutation^36, 38^ via HDR confers dominant cellular resistance to rapamycin in primary T cells.

We then devised the strategy to simultaneously target a therapeutic transgene and introduce the F2108L mutation to the *MTOR* locus (**Fig. 3a**). After targeted integration, *MTOR*-F2108L is expressed on the forward DNA strand while the moderately active human *PGK1* promoter drives the expression of a therapeutic transgene of interest on the reverse DNA strand (**Fig. 3a**). As a first test, we targeted an expression cassette encoding the fluorescent protein mScarlet-I to the *MTOR* locus in CD3^+^ T cells using linear dsDNA donors^47^. Rapamycin selection reproducibly doubled targeted integration frequency and we observed up to 31% of cells expressing the nested reporter transgene (**Supplementary Fig. 5**). Similarly, we achieved 11% knock-in efficacy of a CD19-CAR-2A-EGFP gene cassette (4.6 kb) following treatment for 8 days with rapamycin (**Supplementary Fig. 6**). While encouraging, the use of large linear dsDNA donor caused toxicity, limited proliferation, and was generally prohibitive for highly efficient gene targeting^47^. In addition, these experiments were performed with CD3^+^ T cells retained in leukoreduction system (LRS) chambers that are typically discarded by blood banks. We found that these cells have higher death rates and limited growth potential under our genome editing conditions in contrast to cells isolated from whole blood donations. Nevertheless, taken together, these data illustrate that rapamycin resistant primary T cells can be engineered to express transgenes in a single homology-directed step using CRISPR-Cas9.

**Figure 3.**
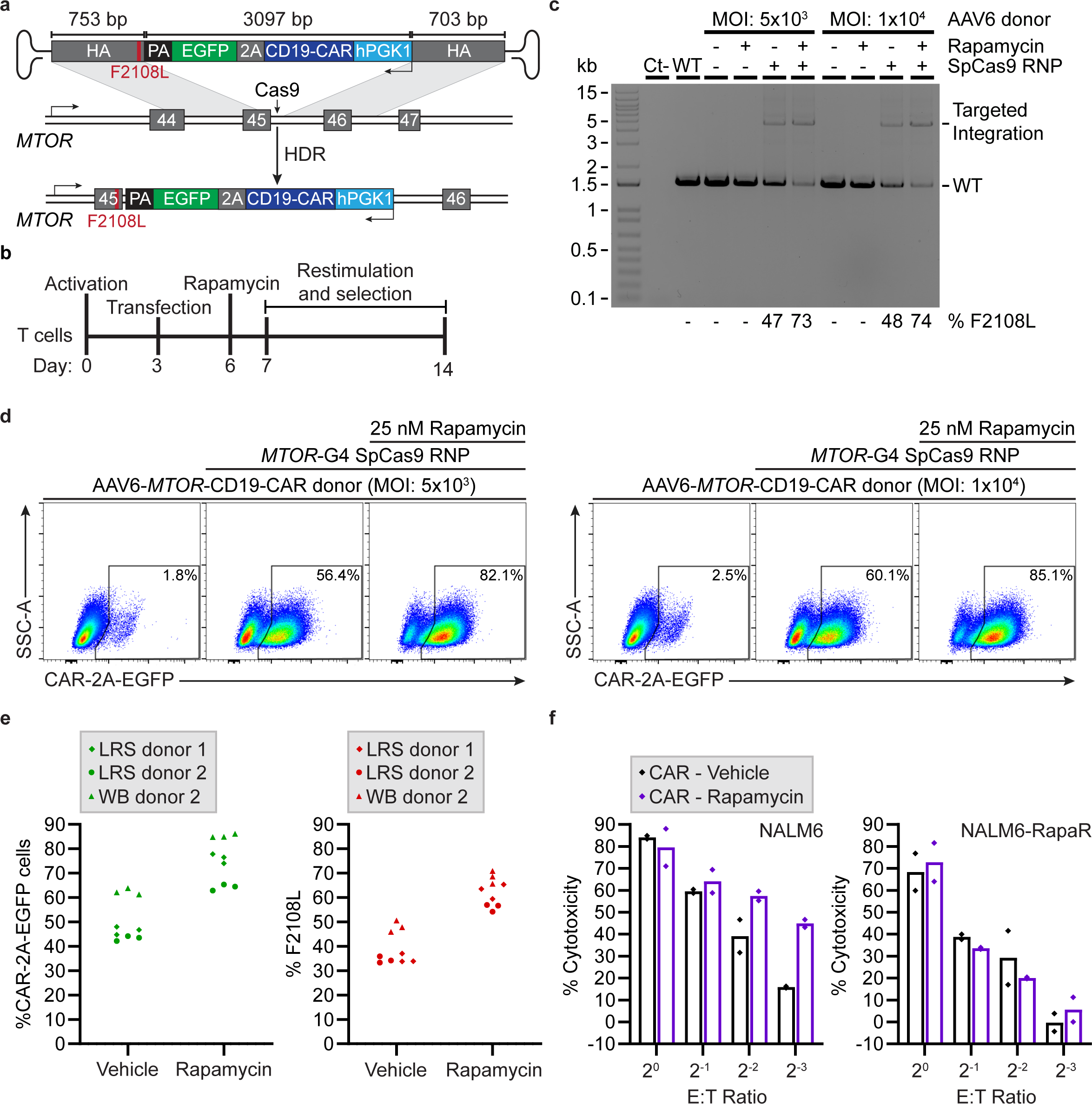
Targeting a CAR to the *MTOR* locus allows the enrichment of rapamycin-resistant CAR-T cells. (**a**) Schematic representation of CD19-CAR-2A-EGFP targeting to the reverse DNA strand of the *MTOR* locus using an AAV6 donor. The F2108L mutation is introduced via the left homology arm (HA) and transgene expression is driven by a human *PGK1* promoter. (**b**) Representative timeline for nucleofection and *ex vivo* rapamycin selection. (**c**) Out-Out PCR for CD19-CAR-2A-EGFP knock-in detection at *MTOR*. Whole blood (WB) primary CD3^+^ T cells from WB donor 1 (WB 1) were transfected with a SpCas9-G4 RNP and transduced with an AAV6 vector with the indicated multiplicity of infection (MOI). T cells were treated (rapamycin) or not (vehicle) with 25 nM rapamycin 3 days post-transfection for 8 days. The percentage of edited alleles was determined by TIDER from Sanger sequencing. *n* = 1 experiment. (**d**) Same as in (**c**), but FACS-based quantification of targeted CD19-CAR-2A-EGFP integration. (**e**) Same as in (**c,d**) with a MOI of 5×10^3^ and primary CD3^+^ T cells isolated from WB or a leukocyte reduction system (LRS) from three additional healthy donors. *n* = 3 independent biological replicates performed in triplicate at different times with CD3^+^ T cells from the indicated healthy donor. Each data point represents a technical replicate. (**e**) Luciferase-based cytotoxicity assay. Following rapamycin selection, CD19-CAR-T cells were incubated with the indicated effector to target (E:T) ratio with NALM6 cells stably expressing firefly luciferase (FLUC) and 25 nM rapamycin. Rapamycin-resistant NALM6 cells harboring the *MTOR*-F2108L mutation (NALM6-RapaR) were used as a control to analyze the combinatorial impact of rapamycin. Luminescence was measured after 18 hours of incubation. *n* = 2 independent biological replicates performed at different times with CD3^+^ T cells from two different healthy donors (WB donor 1 and LRS donor 2). Each data point represents the average of three technical replicates. *hPGK1*, human phosphoglycerate kinase 1 promoter. PA, polyadenylation signal. HA, homology arm.

### One-step generation of rapamycin resistant CAR-T cells

To improve targeting efficacy, we delivered the CD19-CAR-2A-EGFP donor using an adeno-associated virus 6 (AAV6) vector^10, 11, 20^ (**Fig. 3a**). Following Cas9 RNP nucleofection, CD3^+^ T cells isolated from whole blood (Donor ID: WB 1) were transduced with the AAV6 vector in a small volume for 8 hours and then transferred to the culture medium for three days before rapamycin treatment. Cells were reactivated four days post-transfection and expanded for an additional 7 days before quantification of transgene-expressing cells (**Fig. 3b**). We observed a substantial increase in editing efficacy both pre and post 8 days of selection as determined by out-out PCR and Sanger sequencing in the bulk populations (**Fig. 3c**). Yields of CAR-2A-EGFP^+^ cells reached 82% and 85% at MOIs of 5×10^3^ and 1×10^4^, respectively as determined by FACS-based analysis (**Fig. 3d**). These results were reproduced with CD3^+^ T cells isolated from a second whole blood donor (Donor ID: WB2) and two LRS donors (Donor ID: LRS1, LRS2) (**Fig. 3e**). In total, cells from four distinct healthy donors generated similar outcomes, albeit with lower efficiency when using cells isolated from LRS chambers (**Fig. 3e**).

To test for functionality, we performed luciferase-based cytotoxicity assays with selected CAR-T effector cells and CD19^+^ NALM6 target cells stably expressing firefly luciferase and RFP. We also generated a rapamycin-resistant NALM6 cell line (NALM6-RapaR) by installing the *MTOR*-F2108L mutation to assess the combinational impact of rapamycin. We observed high levels of cytotoxicity with and without rapamycin at the highest effector to target ratios (**Fig. 3f**). At lower effector to target ratios, higher levels of cytotoxicity were observed in the presence of rapamycin, confirming that these CAR-T cells can be used in combination with the mTOR inhibitor to target cancer cells (**Fig. 3f**). This combinatorial effect was not observed with NALM6-RapaR cells (**Fig. 3f**). These data indicate that immunosuppressive doses of rapamycin can be used in combination with these CAR-T cells to increase antitumor activity *in vitro*.

Rapamycin has previously been shown to accelerate the memory T-cell differentiation program and may impact the phenotype of treated cells^49, 50^. To begin to probe the impact of rapamycin treatment *ex vivo*, we analyzed differentiation and exhaustion markers and performed multi-color flow cytometry panels after 8 and 15 days of treatment using CD3^+^ T cells isolated from whole blood of two additional healthy donors (ID: WB3, WB4) (**Supplementary Fig. 7**). In these experiments, editing rates remained high (∼80% CAR-2A-EGFP^+^ cells) and we observed a minimal impact of rapamycin treatment on the expression of differentiation and exhaustion markers, as determined by CD4, CD8, CD45RO, CCR7, and PD1 staining, on both rapamycin-sensitive and rapamycin-resistant T cells (**Supplementary Fig. 7**). Rapamycin-resistant CAR-2A-EGFP^+^ populations displayed a mix of central memory (CCR7^+^/CD45RO^+^) and effector memory (CCR7^-^/CD45RO^+^) T cells after 8 days of treatment with rapamycin, while most cells displayed an effector memory (CCR7^-^/CD45RO^+^) phenotype after 15 days of treatment (**Supplementary Fig. 7**). Overall, rapamycin treatment (25 nM) had little impact on T cells’ differentiation in our *ex vivo* expansion protocol, and our 14 days expansion timeline (8 days of rapamycin treatment) yields a mix of central memory and effector memory CAR-T cells.

### Combinatorial CAR-T cell therapy with rapamycin slows leukemia progression *in vivo*

We then established a xenograft leukemia model using NOD/SCID/IL-2Rγ-null (NSG) male and female mice injected with NALM6 cells stably expressing RFP and firefly luciferase. NALM6 cells were injected intravenously, and rapamycin treatment (4 mg/kg, daily) started three days later. Rapamycin-resistant CD19-CAR-2A-EGFP^+^-CAR T cells (Donor ID: WB2, see **Fig. 3e**), which underwent *ex vivo* selection with 25 nM rapamycin for eight days (see **Fig. 3a,b)**, were administered intravenously four days after tumor inoculation (**Fig. 4a)**. In total, these CD19-CAR T cells where expanded *ex vivo* for 14 days and stimulated twice before being injected in mice (see **Fig. 3b)**. As a negative control, we used CD3^+^ T cells untransduced (UT) with AAV6 donors but having undergone nucleofection with the Cas9-*MTOR*-G4 RNP. Tumor engraftment and progression were assessed via bioluminescence imaging twice per week for seven weeks. While rapid tumor progression was observed with UT T cells without treatment, rapamycin slowed tumor progression and conferred an additional week of survival (**Fig. 4b,c)**. Of importance, rapamycin-resistant CD19-CAR-T cells selected *ex vivo* could efficiently target NALM6 cells *in vivo* in the presence or absence of rapamycin (**Fig. 4b,c)**. Combinatorial rapamycin administration with rapamycin-resistant CAR-T cells further slowed tumor progression and allowed the mice to survive for up to seven weeks post-tumor inoculation (**Fig. 4b,c)**. While we observed antitumor control during the first weeks, tumors eventually rebounded and killed all mice (**Fig. 4b,c)**. Our results corroborate previous findings demonstrating the limited persistence of second-generation CARs harboring a CD28 costimulation domain compared to 4-1BB^51–53^. Indeed, the second-generation CD19-CAR architecture used during this study^54^ contains a CD28 transmembrane domain, which can form an heterodimer with CD28^55^, and a CD28 costimulation domain that has been linked to lower persistence compared to 4-1BB^6, 53, 56^. Nevertheless, our results confirmed that our targeting strategy can create functional rapamycin-resistant CAR-T cells that can be used in combination with rapamycin to slow tumor progression *in vivo*, offering an opportunity to target cancer cells’ metabolism to facilitate tumor eradication.

**Figure 4.**
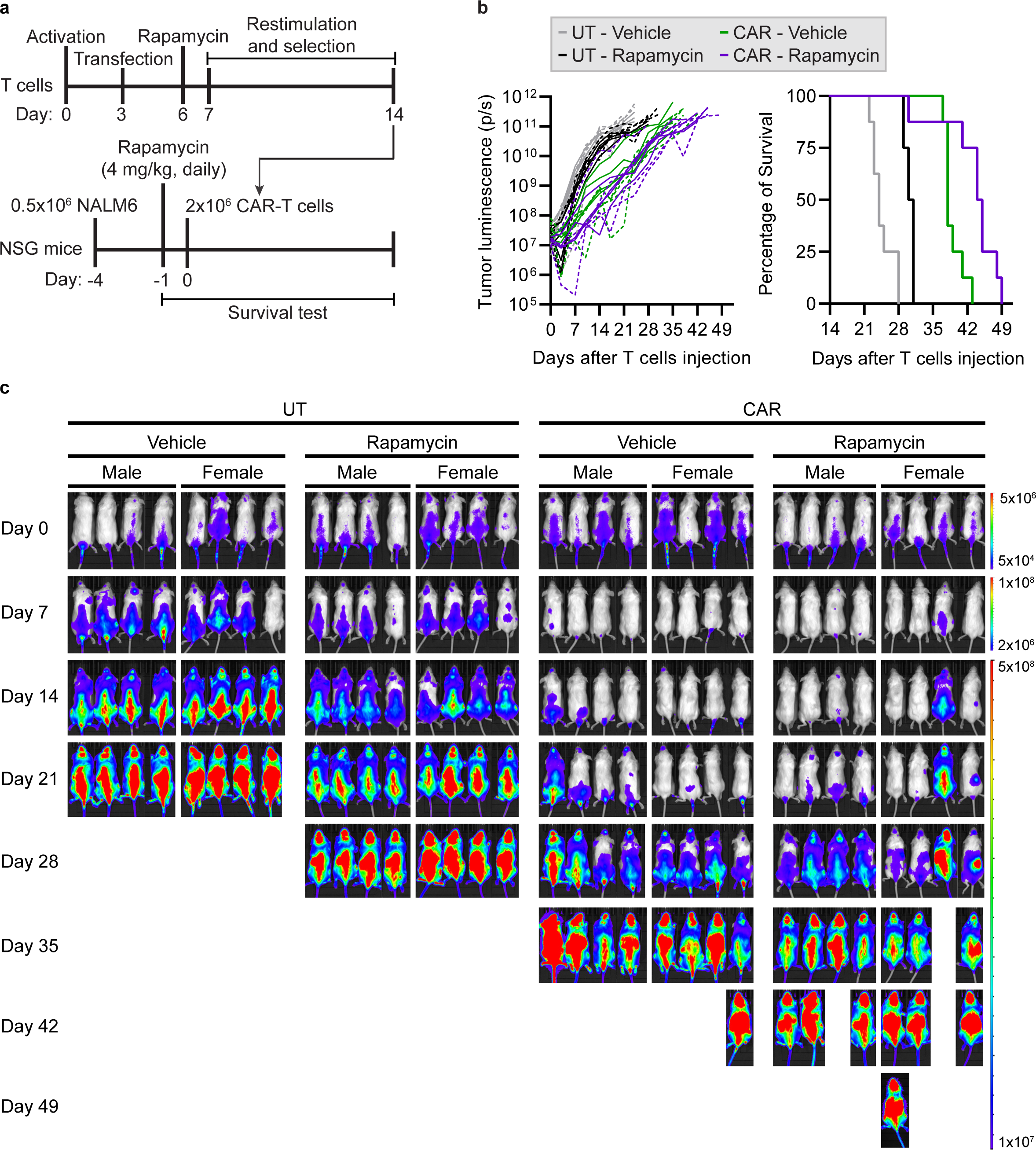
Combinatorial CAR-T cell therapy with rapamycin slows leukemia progression *in vivo*. (**a**) Timeline and experimental setup for the pre-B acute lymphoblastic leukemia xenograft mouse model. Male and female NSG mice were challenged with 0.5 x 10^5^ NALM6-RFP-FLUC cells and daily rapamycin treatment (4 mg/kg) started three days later. Four days after tumor inoculation, 2 x 10^6^ untransduced (UT) T cells or CD19-CAR-T cells were injected (Donor ID WB2, see Fig. 3e). (**b**) Bioluminescence (BLI) quantification and Kaplan-Meier survival plot. Leukemia engraftment, bio-distribution, and tumor progression were assessed by BLI imaging two times per week. The radiance (photons/s) of the regions of interest (ROIs) corresponds to the area containing the whole back side of the body. Radiance quantifications are illustrated with full and dotted lines for male and female mice, respectively. *n* = 8 mice per group. (**c**) BLI and tumor bio-distribution of all mice from (**b**) over seven weeks. The color barcode represents the radiance scale (photons/s/cm^2^/sr).

### Pharmacological control of CAR-T cell activity using rapamycin

Using intracellular^32^ or extracellular^31^ rapamycin-regulated dimerization domains to pharmacologically control chimeric antigen receptor assembly is a promising approach for developing safer CAR-T cell therapies^57^. However, these systems rely on the non-immunosuppressive rapamycin analog AP21967 which was primarily developed as a tool for chemical biology and do not have suitable pharmacokinetic properties for clinical use^32^, or on non-immunosuppressive doses of rapamycin, which limit their clinical potential. Thus, we tested whether we could target a dimerizing agent-regulated immunoreceptor complex (DARIC)^31^, which can be toggled between on and off states using rapamycin, to the *MTOR* locus.

Building on previous studies^10, 31^, we sought to test whether multiplexed targeting of separate DARIC components could be performed at the *MTOR* and *TRAC* loci simultaneously (**Fig. 5a**). Taking advantage of the dynamic regulation of CAR expression at the *TRAC* locus^10^, which averts tonic CAR signaling, we targeted a CD22 recognition domain (DARIC 2) and a 41BB-CD3z signaling domain (DARIC 3) linked by a P2A self-cleaving peptide to the *TRAC* locus (**Fig. 5a**). In parallel, we targeted an additional CD19 recognition domain (DARIC 1) to the *MTOR* locus to confer dominant cellular resistance to rapamycin while providing an additional antigen specificity to potentially overcome antigen escape^6, 7, 56, 58, 59^. Multiplexed targeted integration of both DARIC cassettes was observed at *MTOR* and *TRAC*, as determined by out-out PCR (**Fig. 5b**). An average of 58% of alleles harboring the *MTOR*-F2108L was observed, confirming that multiplexed genome editing did not decrease targeted transgene integration at *MTOR* (**Fig. 5b**). We performed FACS-based cytotoxicity assays using unselected CAR-T cells in the presence or absence of 25 nM rapamycin with DARIC-T cells that were resistant (RapaR-CD19-CD22-DARIC) or not (CD22-DARIC) to rapamycin. Strikingly, cytotoxicity was only observed with rapamycin-resistant CD19-CD22-DARIC-T cells in the presence of rapamycin, confirming DARIC immunoreceptor dimerization only in the presence of the drug (**Fig. 5c**). As observed above, rapamycin inhibited NALM6 cell growth to some extent, but not NALM6-RapaR which harbors the *MTOR*-F2108L mutation (**Fig. 5c**). Rapamycin-sensitive CD22-DARIC-T cells, where the *MTOR*-F2108L-DARIC-1 AAV6 donor was omitted during transduction, could not target cancer cells in the presence of an immunosuppressive dose of 25 nM rapamycin (**Fig. 5c**). We note that the CD22 scFv construction (DARIC 2) we engineered has never been tested for functionality by itself, thus the absence of cytotoxicity of CD22-DARIC cells in presence of rapamycin should be interpreted with caution.

**Figure 5.**
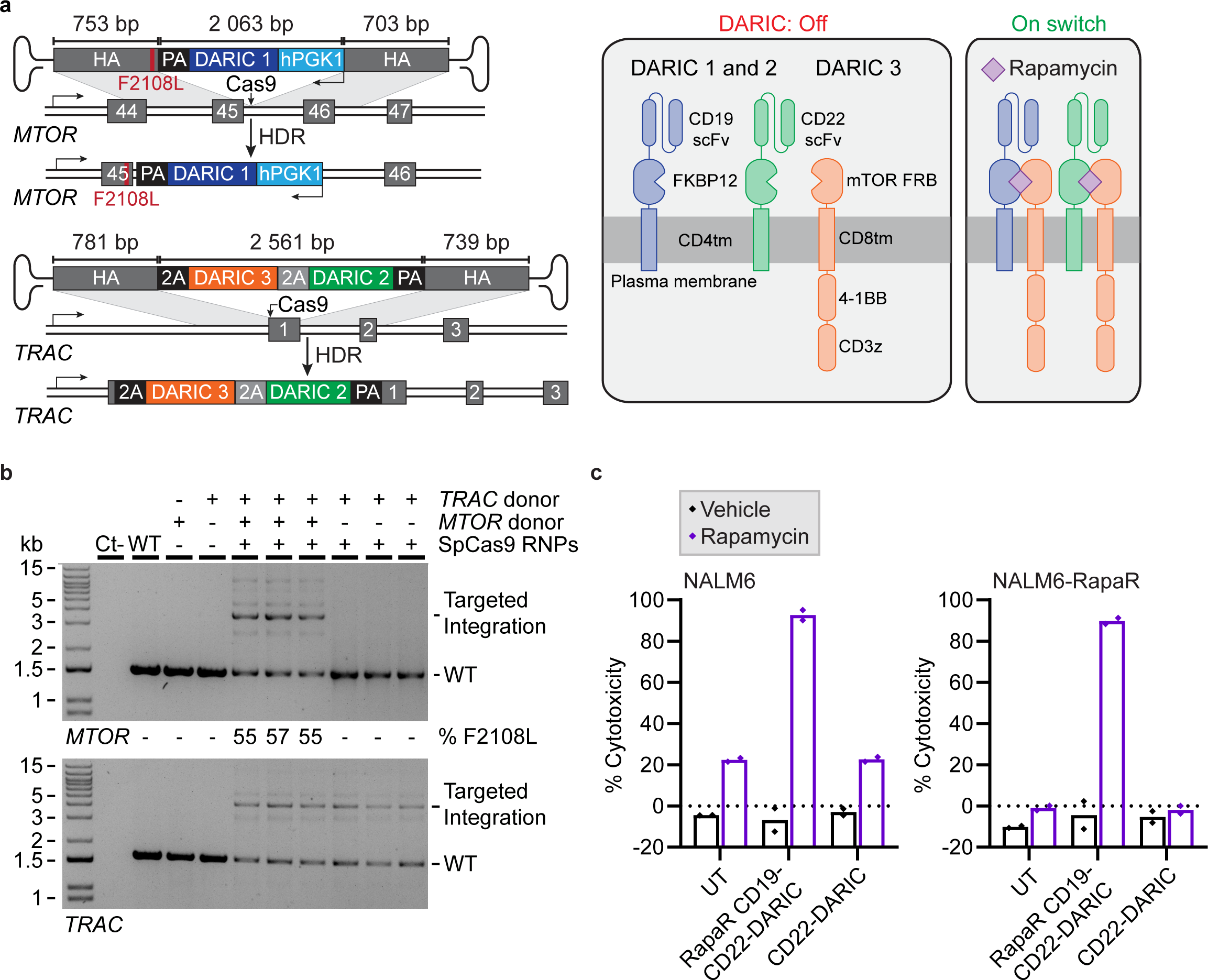
Multiplexed DARIC targeting to the *MTOR* and *TRAC* loci allows pharmacological control of CAR-T cells activity. (**a**) Schematic representation of DARIC-1 targeting to the reverse DNA strand of the *MTOR* locus, and DARIC-2 and 3 targeting to the *TRAC* locus using AAV6 donors. (**b**) Out-Out PCR for DARIC-1 knock-in detection at *MTOR* and DARIC-2 and 3 knock-in at *TRAC*. Whole blood (WB) primary CD3^+^ T cells were transfected with a SpCas9-G4 RNP and transduced with AAV6 vectors with a multiplicity of infection (MOI) of 5×10^3^ for each AAV6. Genomic DNA was harvested 3 days post-transfection. The percentage of edited alleles was determined by TIDER from Sanger sequencing. Representative images are from one of two independent biological replicates. (**c**) FACS-based cytotoxicity assay. CD3^+^ T cells were electroporated with *MTOR*/*TRAC* RNPs and transduced with both *MTOR*/*TRAC* AAV6 donors to generate rapamycin-resistant (RapaR) CD19-CD22-DARIC-T cells. For rapamycin-sensitive CD22-DARIC-T cells, the *MTOR* AAV6 donor was omitted. Three days post-nucleofection, DARIC-T cells were incubated in the presence (rapamycin) or absence (vehicle) of 25 nM rapamycin with NALM6 cells stably expressing RFP and K562-RapaR stably expressing EBFP with an effector to target ratio of 1:1. Rapamycin-resistant NALM6 cells harboring the *MTOR*-F2108L mutation (NALM6-RapaR) were used as a control to analyze the combinatorial impact of rapamycin. The percentage of cytotoxicity was measured by FACS after 72 hours of incubation. *n* = 2 independent biological replicates. Each data point represents the average of three technical replicates.

Using our established xenograft leukemia model, we then tested whether multiplexed DARIC targeting could be used to activate DARIC-T cells *in vivo* even when using an immunosuppressive dose of rapamycin. We targeted DARIC components to the *MTOR* and *TRAC* loci (see **Fig. 5a**) and administered DARIC-T cells to NSG mice three days post-transfection. This time, no prior *ex vivo* expansion and selection was performed before injecting T cells into mice (**Fig. 6a**). No anti-tumor activity was detected in mice treated with DARIC-T cells in the absence of rapamycin, confirming the absence of extracellular receptor dimerization without pharmacological induction (**Fig. 6b,c**). Strikingly, complete tumor control was observed during the first three weeks in all mice treated with rapamycin and DARIC-T cells (**Fig. 6b,c**). While maximal tumor burden was observed in all mice that were not treated with rapamycin three weeks after T cells injection, mice treated with DARIC-T cells in combination with rapamycin survived up to eleven weeks (**Fig. 6b,c and Supplementary Fig. 8**). Of relevance, we observed a prolonged anti-tumor control in female mice (**Fig. 6b,c and Supplementary Fig. 8**). While the impact of sex as a biological variable warrants further investigation, we note that the female NSG mice weighed less on average (∼3-5 grams difference) than their male counterparts at the start of the experiment (see MS source data). Thus, the female mice received both a larger tumor burden and a larger dose of DARIC-T cells. Interestingly, it has recently been reported in a retrospective study of clinical trials for CD19 CAR-T cell therapy that the response and survival are significantly superior in female compared to male patients^60^. Altogether, this multiplexed strategy offers the opportunity to use immunosuppressive doses of rapamycin in combination with DARIC-T cells to decrease tumor cell growth and to pharmacologically fine-tune CAR-T cells’ activity.

**Figure 6.**
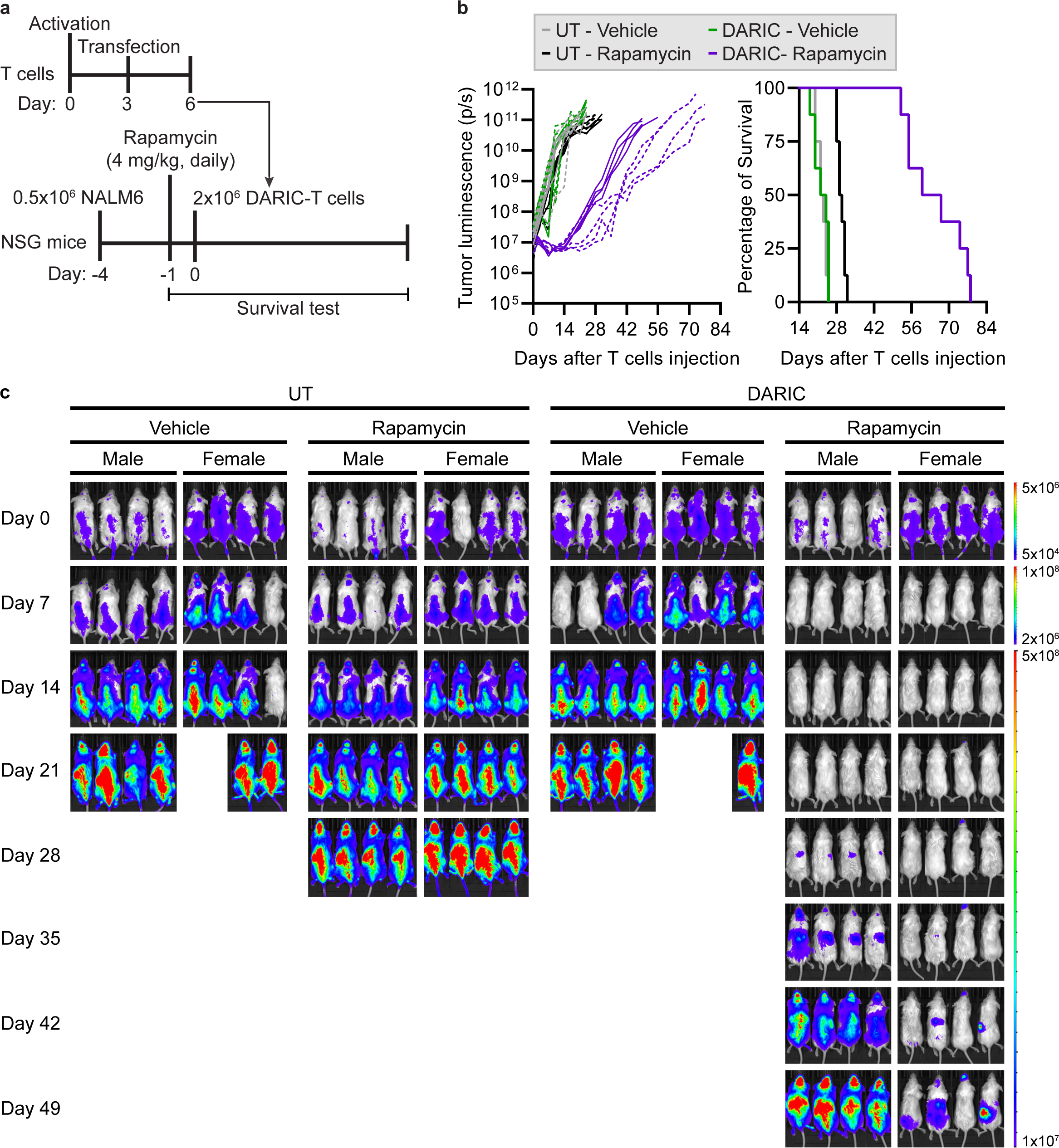
Multiplexed DARIC targeting allows pharmacological activation *in vivo* with an immunosuppressive dose of rapamycin. (**a**) Timeline and experimental setup for the pre-B acute lymphoblastic leukemia xenograft mouse model. Male and female NSG mice were challenged with 0.5 x 10^5^ NALM6-RFP-FLUC cells and daily rapamycin treatment (4 mg/kg) started three days later. Four days after tumor inoculation, 2 x 10^6^ untransduced (UT) T cells or DARIC-T cells were injected. (**b**) Bioluminescence (BLI) quantification and Kaplan-Meier survival plot. Leukemia engraftment, bio-distribution, and tumor progression were assessed by BLI imaging two times per week. The radiance (photons/s) of the regions of interest (ROIs) corresponds to the area containing the whole back side of the body. Radiance quantifications are illustrated with full and dotted lines for male and female mice, respectively. *n* = 8 mice per group. (**c**) BLI and tumor bio-distribution of all mice from (**b**) over seven weeks (See **Supplementary Fig. 8b** for tumor bio-distribution over 11 weeks). The color barcode represents the radiance scale (photons/s/cm^2^/sr).

## DISCUSSION

Engineering primary T cells with programmable nucleases for cell therapy applications remains challenging. In this work, we describe a marker-free selection approach to simultaneously target a therapeutic transgene of interest and a point mutation to the *MTOR* locus to generate dominant cellular resistance to rapamycin. We show that mTORC1 signaling remained functional and cells grew robustly in the presence of rapamycin after targeted transgene integration. Our results further demonstrate that introns provide an additional non-coding location for targeted transgene integration via intron nesting^34, 35^. Taking advantage of this naturally occurring genomic architecture allows the co-modification of an endogenous gene during the integration of a therapeutic transgene of interest in a single gene editing event. However, this strategy is limited to moderately active promoters, such as the human *PGK1* promoter used in this study, since transcriptional interference between the overlapping genes^35^ would be detrimental for mTORC1 signaling, and consequently, CAR-T cell activation and expansion^21, 30^. This could be prohibitive for applications where high levels of transgene expression are needed to reach a therapeutic threshold.

The metabolic competition between cancer and T cells has previously been shown to drive cancer progression^61^, and metabolic interventions could be used in combination to improve clinical outcomes^62^. Hence, targeting cancer metabolism via mTORC1 inhibition could potentially slow tumor progression and facilitate complete tumor eradication. Taken together, our findings suggest that rapamycin could be administered to enrich CRISPR-engineered cells *in vivo* while providing a fitness advantage against tumor cells. Of note, since double editing events at two different loci are not statistically independent, rapamycin co-selection could also be performed to enrich a second modification of interest^34, 63^.

Our cytotoxicity assays corroborate previous findings regarding the combinatorial impact of rapamycin with *piggyBac*-modified rapamycin-resistant CAR-T cells^27^. Nonetheless, our CRISPR-based strategy offers advantages over the use of a two *piggyBac* transposon vectors system to target a CAR and a large rapamycin-resistant mTOR transgene cassette (≈7.65kb)^27^ to random genomic locations^27–29^.

While we focused on the standard CD19^+^ NALM6 acute lymphoblastic leukemia model during this study, combinatorial CAR-T cell therapy with rapamycin could potentially improve antitumor activity against solid tumors displaying hyperactive mTOR signaling^36, 64, 65^. One key advantage of our gene targeting approach is the opportunity to use dimerizing agent-regulated immunoreceptor complex (DARIC) switchable between on and off states using rapamycin to develop safer CAR-T cells therapies^31, 32, 57^. One limitation of the previously reported systems is the use of non-immunosuppressive rapamycin dosing^31^ to prevent the inhibition of CAR-T cells’ activation, which can be overcome using rapamycin-resistant DARIC-T cells. Our system could be used to fine-tune CAR-T cells activity with rapamycin pulsing to prevent tonic CAR signaling, and consequently, exhaustion^31, 66^. In line with this, transient rest induced by a drug-regulatable CAR or the tyrosine kinase inhibitor dasatinib has been shown to restore functionality in exhausted CAR-T cells through epigenetic remodeling^67–69^. Most importantly, rapamycin-regulatable CAR architectures could be used as safety switches to dampen toxicity and severe adverse events associated with cytokine release syndrome^31, 67–71^.

During this study, we also identified new rapamycin resistance mutations via saturation prime editing that displayed intermediate levels of mTORC1 signaling in the presence of rapamycin. These additional mutations offer interesting opportunities to investigate the potential of pharmacologically fine-tuning mTORC1 activity for cell therapy applications. mTOR plays key roles in T cell fate decisions and differentiation^30, 49^. Reducing mTORC1 signaling has been shown to trigger the formation of stem cell-like memory T cells and also enhance CAR-T cell antitumor activity^50, 72^. In addition, it was shown that transient mTOR inhibition with rapamycin rescues 4-1BB CAR-Tregs from tonic signal-induced dysfunction^73^. Considering that mTORC1 signaling acts as a metabolic rheostat^24^, fine-tuning this pathway could potentially have a positive impact on CAR-T cells persistence, and consequently, CAR-T cells antitumor activity. While this hypothesis warrants further investigation, our results demonstrate that engineering the *MTOR* locus with CRISPR allows the marker-free selection of cells with different levels of mTORC1 signaling in the presence of rapamycin.

Altogether, the strategy presented here should facilitate the *ex vivo* and *in vivo* enrichment of CRISPR-engineered cells expressing a therapeutic transgene of interest while offering an opportunity to pharmacologically fine-tune cell signaling, providing a versatile platform for cell therapy applications.

## METHODS

### K562 and NALM6 cell culture and transfection

K562 cells were obtained from the ATCC (CCL-243) and NALM6 cells stably transduced to express a FLUC-T2A-RFP-IRES-Puro reporter gene cassette (Biosettia) were gently provided by Scott McComb (National Research Council of Canada). K562 and NALM6 cells were maintained at 37 °C under 5% CO_2_ in RPMI medium supplemented with 10% FBS, penicillin–streptomycin, and GlutaMAX. Cells were routinely tested for the absence of mycoplasma contamination. Rapamycin (Cayman Chemicals, Cat 53123-88-9) was dissolved at 10 mg/ml in DMSO, working dilutions were prepared in water and stored at −20°C. AZD8055 (STEMCELL Technologies) was dissolved at 10 mM in DMSO, working dilutions were prepared in water and stored at −20°C. DMSO alone was diluted in water and used as vehicle control. Ouabain octahydrate (Sigma) was dissolved at 5 mg/ml in hot water, working dilutions were prepared in water and stored at −20°C. K562 and NALM6 cells (2×10^5^ cells/transfection) were transfected using a Lonza 4D nucleofector™ with a SF nucleofection kit (Lonza) following manufacturer’s recommendations. For prime editing transfections, K562 cells were transfected with 0.75 µg of PE vector, 0.25 µg of epegRNA vector, and 0.1 µg of nick sgRNA vector. NALM6 cells were transfected with 50 pmols of SpCas9 RNP (IDT) and 0.75 µg of dsDNA donor harboring *MTOR*-F2108L and a PAM mutation to generate the NALM6-RapaR cell line. Cells were treated with the indicated concentration of rapamycin 3 days post-nucleofection until all non-resistant cells were eliminated.

### Genome editing vectors and recombinant AAV production

Guide RNAs were designed with CRISPOR^43^ and their sequences are provided in the **Supplementary material** section. When required, DNA sequences for the guides were modified at position 1 to encode a G, owing to the transcription requirement of the human U6 promoter. Guide RNAs were cloned into pX330-U6-Chimeric_BB-CBh-hSpCas9 (a gift from Feng Zhang; Addgene 42230) and their sequences are provided in the **Supplementary material** section. Plasmid donors were cloned into pUC19 with short homology arms (700-800 bp) and all sequences were confirmed by Sanger sequencing. Prime editing (PE) experiments were performed with pCMV-PEmax^74^ (a gift from David Liu; Addgene 174820), and pU6-tevopreq1-GG-acceptor^41^ (a gift from David Liu; Addgene 174038). To repurpose the mTORC1 signaling reporter mVenus-TOSI previously developed for mouse^40^, the N-terminal residues (1-82) of the human *PDCD4* gene were codon optimized using the GenSmart™ codon optimization tool, synthesized as a gBlocks™, and cloned into ATP1A1_T804N_hPGK1_mScarlet-I_Donor (Addgene 173207) upstream of the mScarlet-I-NLS cassette using AflII and NcoI. The CD19-CAR-28z-2A-EGFP gene cassette from pSLCAR-CD19-28z (a gift from Scott McComb: Addgene 135991)^54^ was cloned into the *MTOR* donor vector by Gibson assembly without the 3xFLAG. The CD19-DARIC expression vector was designed in-house based on previously described constructions^31^ and US patent 2012/0266971 A1. The CD22-DARIC construction was designed using a previously described CD22 scFv fragment^59^. DARIC sequences were codon-optimized using GenSmart™ codon optimization tool, synthesized by GenScript, and cloned into the AAV6 vector using Gibson assembly. Vectors used for AAV6 production are available at Addgene and their ID are provided in the **Supplementary material** section. The AAV6 vectors were produced by the vector core facility at the Canadian neurophotonics platform (molecular tools). ITR integrity was assessed following BssHII digestion of the AAV plasmid before production. The virus was resuspended in PBS 320 mM NaCl + 5% D-sorbitol + 0.001% pluronic acid (F-68), aliquoted, and stored at −80°C. The vector yields were 1×10^12^, 8.5×10^12^, and 6.9×10^12^ GC/ml for the *MTOR*-F2108L-CD19-CAR-2A-EGFP, *MTOR*-F2108L_CD19_scFv, and TRAC_T2A_CD22-DARIC donors, respectively.

### Primary human T cells isolation and culture

Primary CD3^+^ human T cells were isolated from anonymized healthy human donors either from fresh whole blood (WB) or leukocyte reduction system (LRS) chambers (Héma-Québec), or from human peripheral blood leukopaks (STEMCELL). EasySep™ Human T cell isolation kit (STEMCELL) or CD3 microbeads (Miltenyi) were used for positive magnetic selection of CD3^+^ T cells from peripheral blood mononuclear cells (PBMCs). Primary CD3^+^ T cells were cultured with Immunocult-XF T cell expansion medium supplemented with 1% penicillin-streptomycin and 300 U/ml IL-2. Cells were thawed and activated for 3 days with Immunocult™ human CD3/CD28/CD2 T cell activator (STEMCELL) before nucleofection.

### Primary human T cells transfection

dsDNA donors were produced as described previously^47^. Briefly, donor sequences were cloned into a cloning vector and PCR amplification (25 cycles) was performed using Kapa-HiFi polymerase (Roche). DNA purification was performed by solid-phase reversible immobilization using AMPure XP (Beckman Coulter) magnetic beads. For ribonucleoprotein complex (RNP) formulation, 100 pmols of sgRNAs (Alt-R, IDT) were mixed with 0.8 µl of 100 mg/ml polyglutamic acid (Sigma) and 50 pmols of SpCas9 nuclease (IDT) as previously described^75^. The RNP mix was incubated at 37°C for 10 minutes and the dsDNA donors were added to the mix and incubated at room temperature for 5 minutes before transfection. Primary CD3^+^ T cells (1×10^6^ cells/transfection) were transfected using a Lonza 4D nucleofector™ (Pulse code EH115) with a P3 nucleofection kit (Lonza) following manufacturer’s recommendations with the indicated DNA concentrations. CD3^+^ T cells were resuspended in 80 µl media without IL-2 directly after nucleofection and incubated at 37°C for 15 minutes before transfer into culture media plates at 1×10^6^ cells/ml. For AAV6 transduction, CD3^+^ T cells were resuspended in 80 µl media directly after nucleofection and AAV6 vectors were added. Cells were incubated at 37°C for 8 hours at high cell density before transfer into culture media plates. For rapamycin selection, CD3^+^ T cells were treated with or without 25 nM rapamycin on Day 3 post-nucleofection, restimulated with Immunocult™ human CD3/CD28/CD2 T cell activator (STEMCELL) 4 days post-transfection, and expanded for an additional 7 days.

### Sanger sequencing analysis and out-out PCRs

Genomic DNA was extracted with QuickExtract DNA extraction solution (EpiCentre) following manufacturer’s recommendations. Primers used in this study and the PCR product sizes are provided in the **Supplementary material** section. PCR amplifications were performed with 30 cycles of amplification with Phusion polymerase. Sanger sequencing was performed on PCR amplicons to quantify the percentage of edited alleles using TIDE^76^, TIDER^77^, and BEAT^78^. To quantify the percentage of alleles harboring the *MTOR*-F2108L mutation after targeted transgene integration, we performed PCR amplifications to amplify both WT and HDR alleles with a forward primer binding outside the region of homology of our donors, and a reverse primer binding inside the homology arm upstream of the SpCas9 target sites (out-in PCR). Genomic DNA from WT cells and cells transfected or transduced only with the donor were used as negative controls in all experiments. Kapa-HiFi polymerase was used for out-out PCRs. To detect targeted integration, out-out PCRs were performed with primers that bind outside of the homology regions of the plasmid/dsDNA donors. Wild type K562 or CD3^+^ T cells genomic DNA was used as a control for all PCRs.

### Flow cytometry

The percentage of fluorescent cells was quantified using a BD LSRII flow cytometer using FACSdiva v6.1.2 software, and 1×10^5^ cells were analyzed for each condition. For mTORC1 signaling assays, cells were washed with PBS and cultured at least 3 days without rapamycin. Cells were treated with the indicated concentration of rapamycin or AZD8055 for 24 hours before analysis. FACS sorting was performed using a BD FACS Aria Fusion flow cytometer for the saturation prime editing experiment, and 5×10^5^ cells were sorted for each condition.

Multicolor flow cytometry panels were performed using a Cytek Aurora flow cytometer. T cells were stained with fixable viability dye eF506 (Invitrogen Cat 65-0866-18) for 15 minutes at 4°C, then washed and resuspended in Human TruStain FcX blocking solution (BioLegend Cat 2711505) and Brilliant Stain Buffer Plus (BD Biosciences Cat 566385) for 10 minutes at room temperature. After blocking, cells were stained with the following antibody cocktail diluted in flow cytometry staining buffer: mouse anti-human CD3 BV750 (BioLegend Cat 344846), mouse anti-human CD4 AF700 (BioLegend Cat 300526), mouse anti-human CD8 PerCP (BioLegend Cat 301030), mouse anti-human CD279 BV421 (BioLegend Cat 329920), mouse anti-human CCR7 APC-Fire 750 (BioLegend Cat 353246), and mouse anti-human CD45RO PE-Cy7 (Invitrogen Cat 25-0457-42). Flow cytometric data were analyzed using SpectroFlo (Cytek) and FlowJo V10 (Tree Star).

### Saturation prime editing and high-throughput DNA sequencing

The *MTOR*-F2108 saturation epegRNA (tevopreQ1)^41^ vector was designed to install NNK codons and a silent R2109R PAM mutation at *MTOR* exon 45, synthesized as a gBlock™ (IDT), and cloned into pU6-tevopreq1-GG-acceptor^41^ (a gift from David Liu; Addgene 174038). K562 cells stably expressing the mSc-TOSI reporter were transfected with PE3max-epegRNA (tevopreQ1) vectors and treated with 0.5 µM rapamycin starting 3 days post-transfection until all non-resistant cells were eliminated. Following selection, cells were treated for 24 hours with 0.5 µM rapamycin and FACS-sorted for low mSc-TOSI fluorescence intensity to enrich cells with functional mTORC1 signaling in the presence of rapamycin. Genomic DNA was harvested after selection and after FACS sorting for low and high mSc-TOSI fluorescence intensity. For amplicon sequencing, primers containing Illumina forward and reverse adapters were used for a first round of PCR using the Kapa-HiFi polymerase. PCR products were purified using AMPure XP magnetic beads and their quality was evaluated by electrophoresis. A second round of PCR and bead purification was performed for indexing, and amplicons were sequenced on an Illumina MiSeq instrument. Alignment of amplicon sequences to the reference *MTOR* sequence was performed using CRISPResso2^79^, and all possible NNK codons were detected.

### GUIDE-Seq

GUIDE-Seq was performed as previously described^45, 46^, except for improvements of the Next Generation Sequencing (NGS) library preparation procedure reported below. GUIDE-Seq oligos harboring 5’ phosphorylation modification and two phosphorothioate linkages between the last two nucleotides at the 3’ end were synthesized by IDT. Oligos were mixed in 50 mM NaCl, 10 mM Tris-HCl (pH 8.0), 1 mM EDTA, and annealed by heating the solution to 95°C for 10 minutes, followed by gradual cooling on a thermocycler. For K562 cells, 2×10^5^ cells were electroporated with 750 ng of SpCas9-sgRNA vectors and 100 pmol of annealed GUIDE-Seq oligos. For primary CD3^+^ T cells, 1×10^6^ cells were electroporated with 50 pmol SpCas9 RNP and 100 pmol annealed GUIDE-Seq oligos. Genomic DNA was harvested using QIAmp UCP DNA Micro Kit (QIAGEN). GUIDE-Seq oligo tag integration was confirmed using TIDE^76^ and DECODR^80^ webtools from Sanger sequencing. The library preparation method was improved starting from the original GUIDE-Seq procedure^45, 46^. After mechanical fragmentation, end-repair and A-Tailing were directly performed using the NEBNext UltraII DNA kit (New England Biolabs). A new custom universal Y-adapter based on the Illumina TruSeq sequences was designed to include a Unique Molecular Index (UMI) in the sequencing read instead of the index read. This modification enables mixing of GUIDE-Seq libraries with other Illumina library types (amplicon, shotgun, RNASeq, etc.) in the sequencing run by using standard index read lengths. This new adapter was ligated as described in the instructions of the NEBNext UltraII procedure using a final concentration of 133 nM. Ligated DNA was purified using 0.9X volume of AMPure XP beads (Beckman-Coulter), then split in 2 for PCR amplifications with either Minus or Plus primers using Q5 DNA Polymerase (New England Biolabs) with the following cycling conditions: 98°C 30 sec; 10 cycles of 98°C 10 sec, 55°C 30 sec, 72°C 30 sec; 15 cycles of 98°C 10 sec, 65°C 30 sec, 72°C 30 sec; 72°C 2 min; 4°C hold. New primers with Illumina sequence tails were specifically designed for this improved method. The use of Illumina sequences at the Plus/Minus amplification stage eliminates the use of custom primers for sequencing. PCR products were purified with 0.9X AMPure XP beads and quantified using Qubit dsDNA HS Assay (Invitrogen). Then, 10 ng DNA was used for a second PCR amplification with Illumina TruSeq dual-indexing primers (98°C 30 sec; 8 cycles of 98°C 10 sec, 55°C 30 sec, 72°C 30 sec; 72°C 2 min; 4°C hold). The barcoding at the PCR stage enables a much simpler and higher multiplexing capacity. PCR products were purified using 0.85X AMPure XP beads, quantified using Qubit dsDNA HS Assay and ran on a Bioanalyzer High Sensitivity DNA chip (Agilent). Samples were pooled in equimolar amounts and sequenced on 1.4% (35M read block) of a NovaSeq S4 PE150 lane (Illumina) at the Centre d’Expertise et de Services Génome Québec (Montréal, QC). The remaining portion of the sequencing lane contained a mixture of library types from multiple organisms (*M. plutonius*, *P. larvae*, *F. graminearum*, *P. neglectus*, *Colletotrichum* spp., *Sporothrix* spp.).

### Analysis of GUIDE-Seq sequencing

A new pipeline, named guideseq_ibis, was designed to analyze the data from the improved GUIDE-Seq procedure (https://github.com/enormandeau/guideseq_ibis). Using this pipeline, the raw reads were trimmed for quality using trimmomatic (v0.36, min_length 100, crop_length 200). Then, only reads containing the expected alien sequence (maximum hamming distance of 1) and only one copy of the ODN sequence (maximum hamming distance of 6) were retained. For these reads, the Unique Read Identifiers (UMIs) and first eight nucleotides were used to rename the sequence. These tagged reads were then mapped onto the latest GRCh38 human genome assembly with bwa (v0.7.17-r1188, -T 10) and then samtools (v1.12, -S -q 1 -F 4 -F 256 -F 2048). The alignment sam files were then sorted by chromosome name and position. The UMI and eight first nucleotides of the reads and the starting positions of their alignments were used to detect duplicated reads. Namely, only one copy of reads with identical UMIs, eight first nucleotides, and alignment start position were kept. Then, Double Stranded Breaks (DSBs) were identified. For this, alignment sites were scanned for peaks of coverage in decreasing order of depth of coverage and pairs of ODN+ and ODN-were reported until the pairs did not meet the minimum coverage threshold (min_length 100, min_coverage 50, window_size 10, position_error 5, bin_size 10000). To account for small errors in sequencing and mapping, read counts are collected within ‘window_size’ nucleotides around each peak and pairs of ODN sequences were kept if they fell withing ‘position_error’ nucleotides of their relative expected positions. Identified sequences on target and off-target were reported for each sample with their chromosome and position localisation, the identifier and name of any gene they overlapped (using a simplifited annotation table from GRCh38.p14), and metrics about the counts of the ODN+ and ODN-reads. Finally, annotation of the found targets and off-targets was performed to add information about the guide’s name, the distance between the target sequence and the expected guide sequence, as well as a representation of the modified nucleotides along the guide sequence. The chromosomal positions and the targeted gene (were applicable) of all off-targets are provided in the **Supplementary material** section.

### Cytotoxicity assays

Luciferase-based cytotoxic assays were performed as previously described^10, 81^. After rapamycin selection, CD19-CAR-T cells were washed with PBS, and co-cultured in 96-well plates in supplemented RPMI medium with NALM6 cells stably expressing firefly luciferase and RFP at the indicated effector to target ratio (E:T) for 18 hours. Target cells incubated with WT CD3^+^ T cells were used to determine maximal luciferase expression (maximal relative light units; RLU_max_). After coculture, an equal volume of luciferase substrate (Bright-Glo, Promega) was added to each well. Emitted light was detected with a Tecan luminescence plate reader and cell lysis was determined as (1 – (RLU_sample_)/(RLU_max_)) x 100. FACS-based cytotoxicity assays were performed as previously described^31^, except with a prolonged coculture time. T cells were washed with PBS, and cocultured in 12-well plates in supplemented RPMI medium with 50:50 mixtures of rapamycin-resistant K562-EBFP (CD19-negative) cells and NALM6-RFP cells at the indicated effector to target ratio (E:T) for 72 hours. Cell viability was assessed using the fixable viability dye eFluor™ 660 (eBioscience). The ratio of viable NALM6-RFP and K562-EBFP cells following coculture in the presence or absence of 25 nM rapamycin was used to calculate the percentage of cytotoxicity. Target cells incubated with WT CD3^+^ T cells were used to determine maximal NALM6-RFP growth.

### Mouse systemic tumor model

The *in vivo* CAR-T cells experiments were performed according to the *Canadian Guide for the Care and Use of Laboratory Animals*. The Université Laval Animal Care and Use Committee approved the procedure. 8-10 weeks-old NOD/SCID/IL-2Rγ-null (NSG) male and female mice were acquired from Jackson Laboratory. 0.5 x 10^6^ NALM6-RFP-FLUC cells were administered via tail vein injection. Three days later, daily rapamycin treatment (4 mg/kg) or vehicle (filter-sterilized solution of 5% Tween80, 5% PEG-400, 0.7% DMSO) was administered by intraperitoneal (IP) injection. Four days after tumor inoculation, 2 x 10^6^ CAR-T cells (≈86% CAR-2A-EGFP^+^ cells) or untransduced (UT) T cells were administered via tail vein injection. CAR-T cells expanded for 14 days, including 8 days of treatment/selection with 25 nM rapamycin, were used for the *in vivo* experiment. The same protocol was used for the multiplexed DARIC-T cells experiment, but DARIC-T cells were administered three days post-transfection without *ex vivo* expansion/selection. Tumor burden was measured twice per week by bioluminescence imaging using the IVIS LuminaIII Imaging System (PerkinElmer). Mice were anesthetized using isoflurane, and 150 mg/kg D-Luciferin (Goldbio) was administered via IP injection. Data were analyzed using the Living Image software (PerkinElmer). The maximum tumor burden and survival endpoint were defined by hindlimb paralysis or other clinical signs of distress. There were no instances in which this maximum tumor burden was exceeded.

## Supporting information

Supplemental Material (DNA sequences, Oligos, Addgene)

## DATA AVAILABILITY

Source data for the figures are provided as a Source Data file. All raw Sanger sequencing data generated in this study are available on request from the corresponding author [YD].

## CODE AVAILABILITY

The adapted GUIDE-Seq analysis pipeline developed and used in this study is freely available at https://github.com/enormandeau/guideseq_ibis.

## ACKNOWLEDGEMENTS

This study was supported by a grant from Mitacs and by the Fondation du CHU de Québec. JJ. L. is supported by a CIHR Project Grant (PJT407274). BC Cancer Foundation funded B.H.N. S.L. was supported by a Frederick Banting and Charles Best Canada graduate scholarship from CIHR, and a Graduate student scholarship from the Fonds de la Recherche du Québec-Santé (FRQS). G.C. holds a Vanier Canada graduate scholarship from CIHR. Salary support was provided by the FRQS to Y.D. B.B. and G.M. are supported by the “Programme d’appui aux plateformes technologiques stratégiques” from the Ministère de l’Économie, de l’Innovation et de l’Énergie Québec. We thank the staff at the flow cytometry core facility of the CHU de Quebec Research Center for their training and assistance. We thank Marie-Ève Paquet and the vector core facility staff at the Canadian neurophotonics platform for AAV6 production. We also thank Scott McComb (National Research Council) for kindly providing NALM6-RFP-FLUC cells.

## AUTHOR’S CONTRIBUTIONS

Conceptualization, S.L. and Y.D.; methodology, S.L., G.C., V.D., C.G., JP. F., S.V., E.N., G.M., N.D., J.L., B.B., JJ. L., and Y.D.; investigation, S.L., G.C., V.D., C.G., JP. F., S.V., E.N., G.M., N.D., J.L., B.B., JJ. L., and Y.D.; writing original draft, S.L.; writing, review and editing, S.L., and Y.D.; supervision, J.L., B.B., JJ. L., and Y.D.; funding acquisition, J.L., JJ. L., and Y.D. All authors reviewed the manuscript and approved its final version.

