## Supplementary Figures for "Pharmacological control of CAR T cells through CRISPR-driven rapamycin resistance"

[1]Centre Hospitalier Universitaire de Québec Research Center - Université Laval, Québec, QC, G1V 4G2, Canada.
[2]Université Laval Cancer Research Centre, Québec, QC, G1V 0A6, Canada.
[3]Present Address: Division of Hematology/Oncology, Boston Children's Hospital. Department of Pediatric Oncology, Dana-Farber Cancer Institute. Harvard Stem Cell Institute. Broad Institute of MIT and Harvard. Department of Pediatrics, Harvard Medical School, Boston, MA 02115, USA.
[4]Trev and Joyce Deeley Research Centre, BC Cancer, Victoria, BC, V8R 6V5, Canada.
[5]Department of Biochemistry and Microbiology, University of Victoria, Victoria, BC, V8P 5C2, Canada.
[6]Institut de Biologie Intégrative des Systèmes, Université Laval, Québec, QC, G1V 0A6.
[7]Medical Affairs and Innovation, Héma-Québec, Québec, Quebec, Canada

[*]Correspondence:
Yannick Doyon, Ph.D.
Centre de recherche du CHU de Québec – Université Laval
2705, boulevard Laurier, T-3-67
Québec, QC G1V 4G2
CANADA


ORCID 0000-0002-3889-9563

**Supplementary figures:**

1. Identification of mutations conferring rapamycin resistance via saturation prime editing
2. *MTOR*-F2108 mutations display different levels of mTORC1 signaling upon rapamycin treatment.
3. Identification of potential off-target sites in K562 and primary CD3$^+$ T cells.
4. Introducing the F2108L mutation to the *MTOR* locus allows the selection of rapamycin-resistant CD3$^+$ T cells.
5. Targeting a fluorescent reporter to the *MTOR* locus allows the selection of rapamycin-resistant CD3$^+$ T cells.
6. Non-viral targeting of a CAR to the *MTOR* locus allows the selection of rapamycin-resistant CAR-T cells.
7. Rapamycin has minimal impact on T cells' differentiation.
8. Prolonged tumor control in female mice treated with rapamycin and DARIC-T cells.

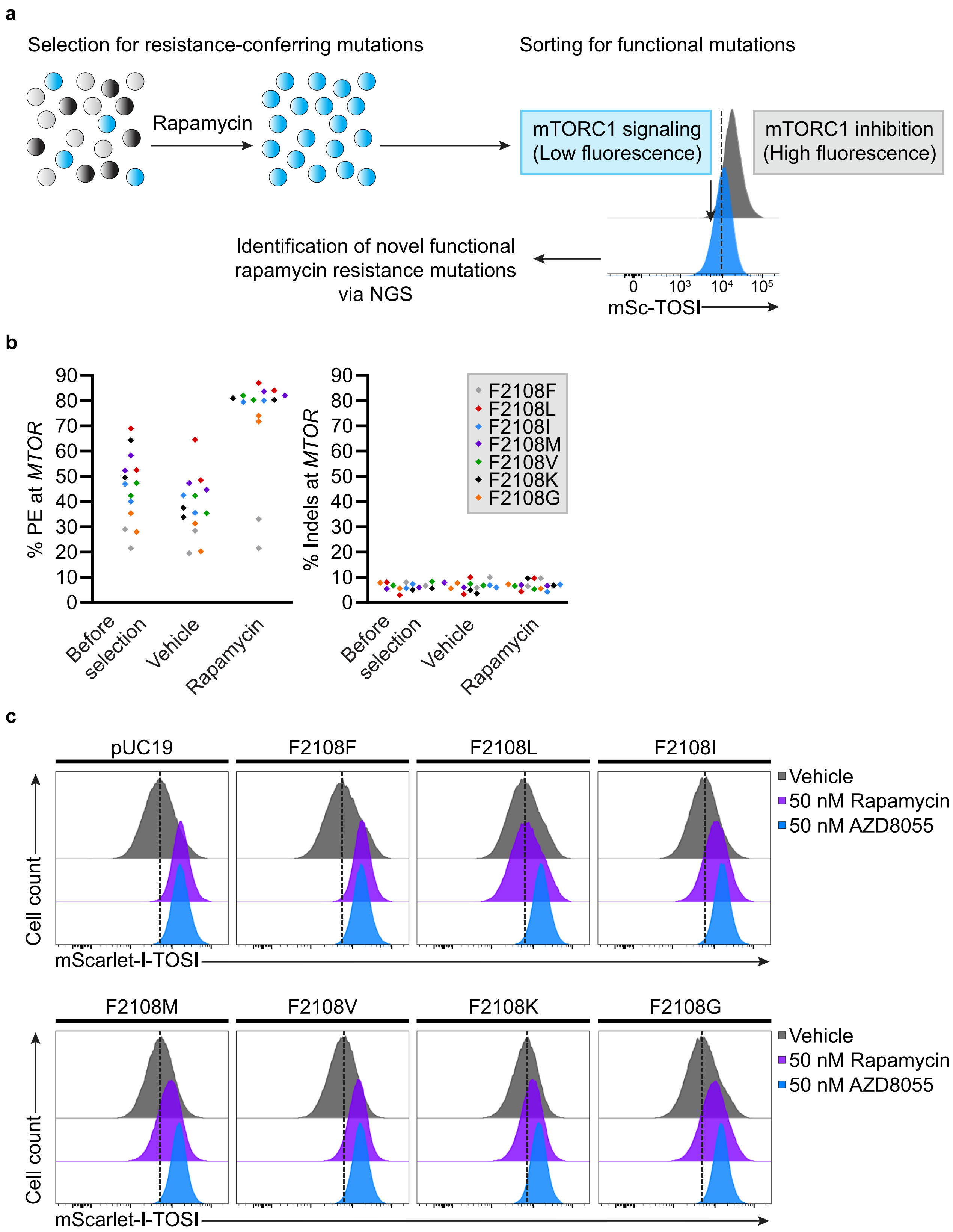


**Figure S1**
**Levesque et al.**

**Supplementary Figure 1 (Related to Fig. 1). Identification of mutations conferring rapamycin resistance via saturation prime editing.** (**a**) Schematic of the saturation prime editing strategy to identify new rapamycin resistance mutations. K562 cells stably expressing the mSc-TOSI reporter are first transfected with PE3 vectors to install all possible amino acid substitutions (NNK codon) at the *MTOR*-F2108 position. Cells are treated with rapamycin until all non-resistant cells are eliminated and FACS sorting is performed to isolate cells displaying functional mTORC1 signaling in the presence of rapamycin. Finally, high-throughput sequencing is performed to identify new functional rapamycin resistance mutations. The representative FACS image is from one of two independent biological replicates performed at different times with equivalent results. Cells were treated with 0.5 µM rapamycin 24 hours before FACS analysis. (**b**) PE and small indels quantification as determined by BEAT and TIDE analysis from Sanger sequencing. K562 cells stably expressing the mSc-TOSI reporter were transfected with PE3max-epegRNA vectors targeting *MTOR*. Genomic DNA was harvested 3 days post-transfection (before selection) and cells were treated (rapamycin) or not (vehicle) with 0.5 µM rapamycin until all non-resistant cells were eliminated. $n$ = 2 independent biological replicates performed at different times. (**c**) Histogram plot of mSc-TOSI intensity in bulk populations of rapamycin-selected cells harboring different *MTOR*-F2108 mutations. Where indicated, cells were treated for 24 hours with 50 nM rapamycin or 50 nM AZD8055 before FACS analysis. Representative images are from one of three independent biological replicates performed at different times with equivalent results (see **Supplementary Fig. 2**).

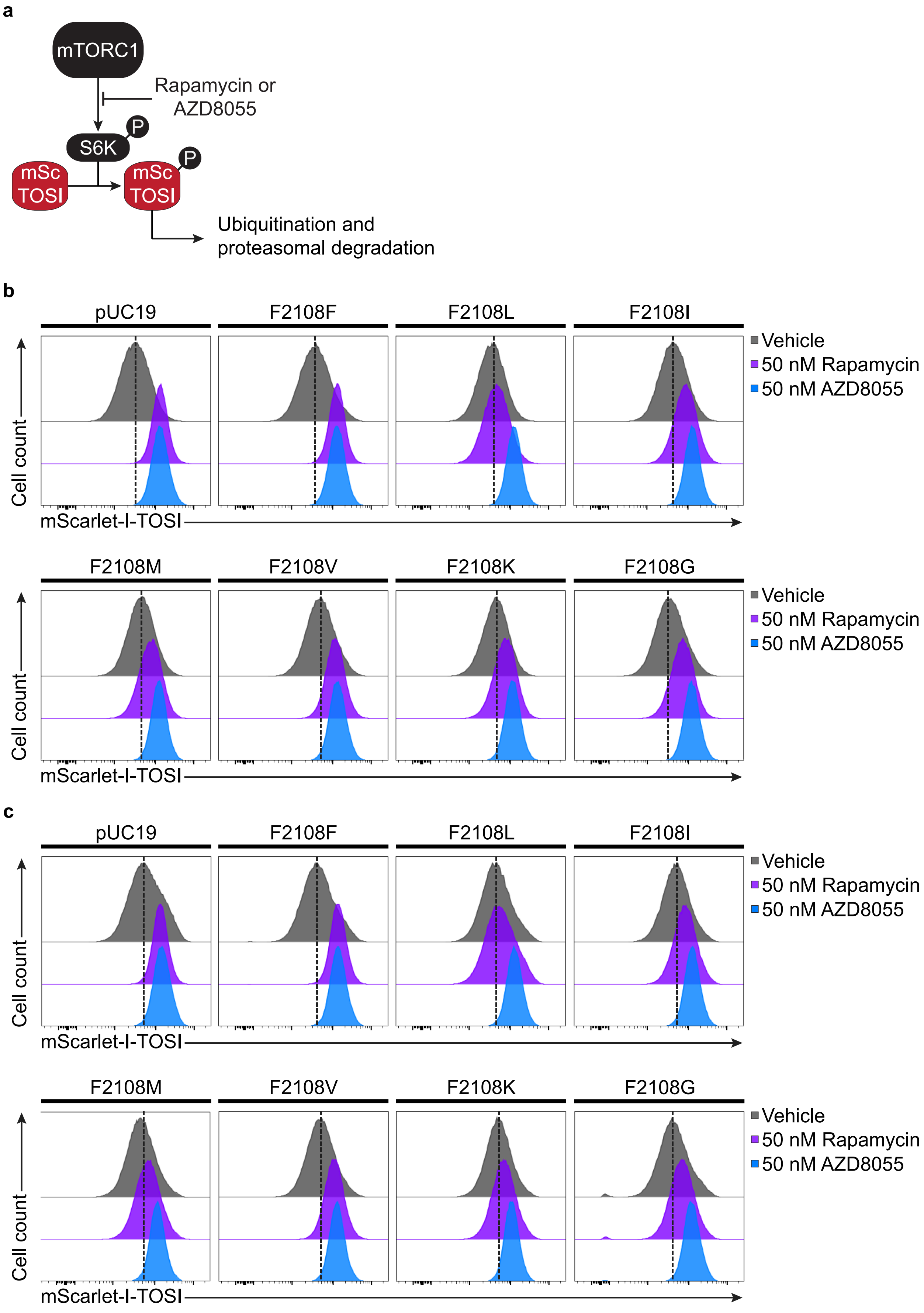


**Figure S2**
**Levesque et al.**

**Supplementary Figure 2 (Related to Fig. 1 and Supplementary Fig. 2). *MTOR*-F2108 mutations display different levels of mTORC1 signaling upon rapamycin treatment.** (**a**) Schematic of mSc-TOSI degradation under mTORC1 signaling. (**b**) Histogram plot of mSc-TOSI intensity in bulk populations of rapamycin-selected cells harboring different *MTOR*-F2108 mutations. Where indicated, cells were treated for 24 hours with 50 nM rapamycin or 50 nM AZD8055 before FACS analysis. As described in **Supplementary Fig. 1**, K562 cells stably expressing the mSc-TOSI reporter were transfected with PE3max-epegRNA vectors targeting *MTOR* and cells were treated with 0.5 µM rapamycin 3 days post-transfection until all non-resistant cells were eliminated. (**c**) Same as in (**b**) with bulk populations of cells from an independent nucleofection. Representative images are from one of three independent biological replicates performed at different times with equivalent results (see **Supplementary Fig. 1**).

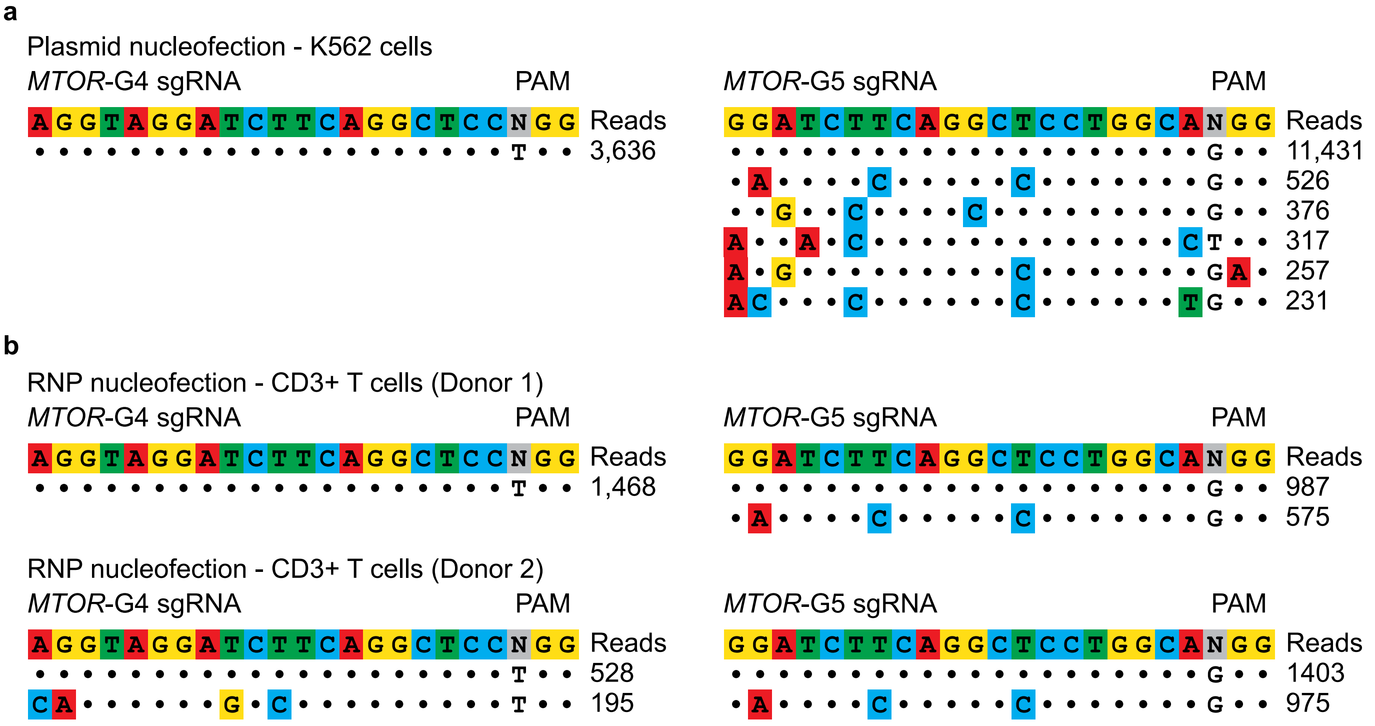


**Figure S3**
**Levesque et al.**

**Supplementary Figure 3 (Related to Fig. 2). Identification of potential off-target sites in K562 and primary CD3$^+$ T cells. (a)** Candidate off-target sites identified in K562 cells using GUIDE-Seq. K562 cells were transfected with a SpCas9-sgRNA-expressing vector and GUIDE-Seq dsDNA tag. Genomic DNA was harvested three days post-nucleofection. Dots represent matches with the intended target sequence, mismatches are colored, and nucleotide bulges are highlighted with a star. GUIDE-Seq read counts are from one experiment. (**b**) Same as (**a**) with primary CD3$^+$ T cells transfected with Cas9 RNPs and GUIDE-Seq dsDNA tag. Genomic DNA was harvested three days post-nucleofection. $n = 2$ independent biological replicates performed with CD3$^+$ T cells from two different donors. Four days post-nucleofection, CD3$^+$ T cells from donor 2 were restimulated and expanded for an additional 7 days before harvesting genomic DNA and the results are illustrated in **Fig. 2**.

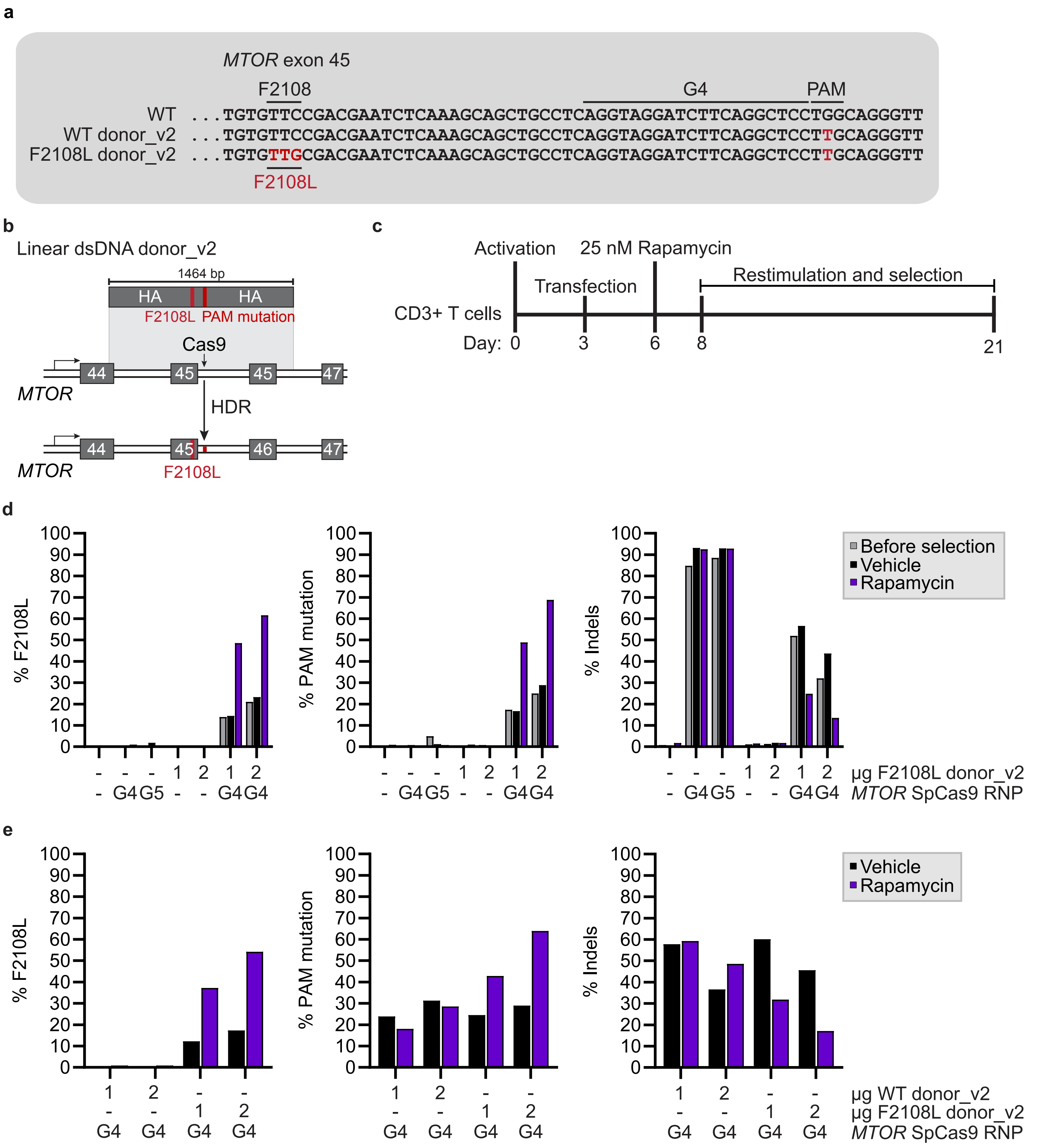


**Figure S4**
**Levesque et al.**

**Supplementary Figure 4. Introducing the F2108L mutation to the *MTOR* locus allows the selection of rapamycin-resistant $CD3^+$ T cells.** (**a**) Schematic representation of *MTOR* intron 45 SpCas9 target sites and the donor sequences used to install the F2108L mutation at *MTOR* exon 45. (**b**) Schematic of the CRISPR-driven installation of the F2108L mutation at *MTOR* using a linear dsDNA donor. (**c**) Timeline for transfection, rapamycin selection, and T cells restimulation. (**d**) HDR quantification as determined by TIDER from Sanger sequencing. $CD3^+$ T cells were transfected with SpCas9-G4 RNP and the indicated amount of linear dsDNA donor. Cells were treated (rapamycin) or not (vehicle) with 25 nM rapamycin 3 days post-transfection for 15 days. $n$ = 1 experiment. (**e**) Independent biological replicate as described in (**d**) but using a WT linear dsDNA donor as a negative control. $n$ = 1 additional experiment.

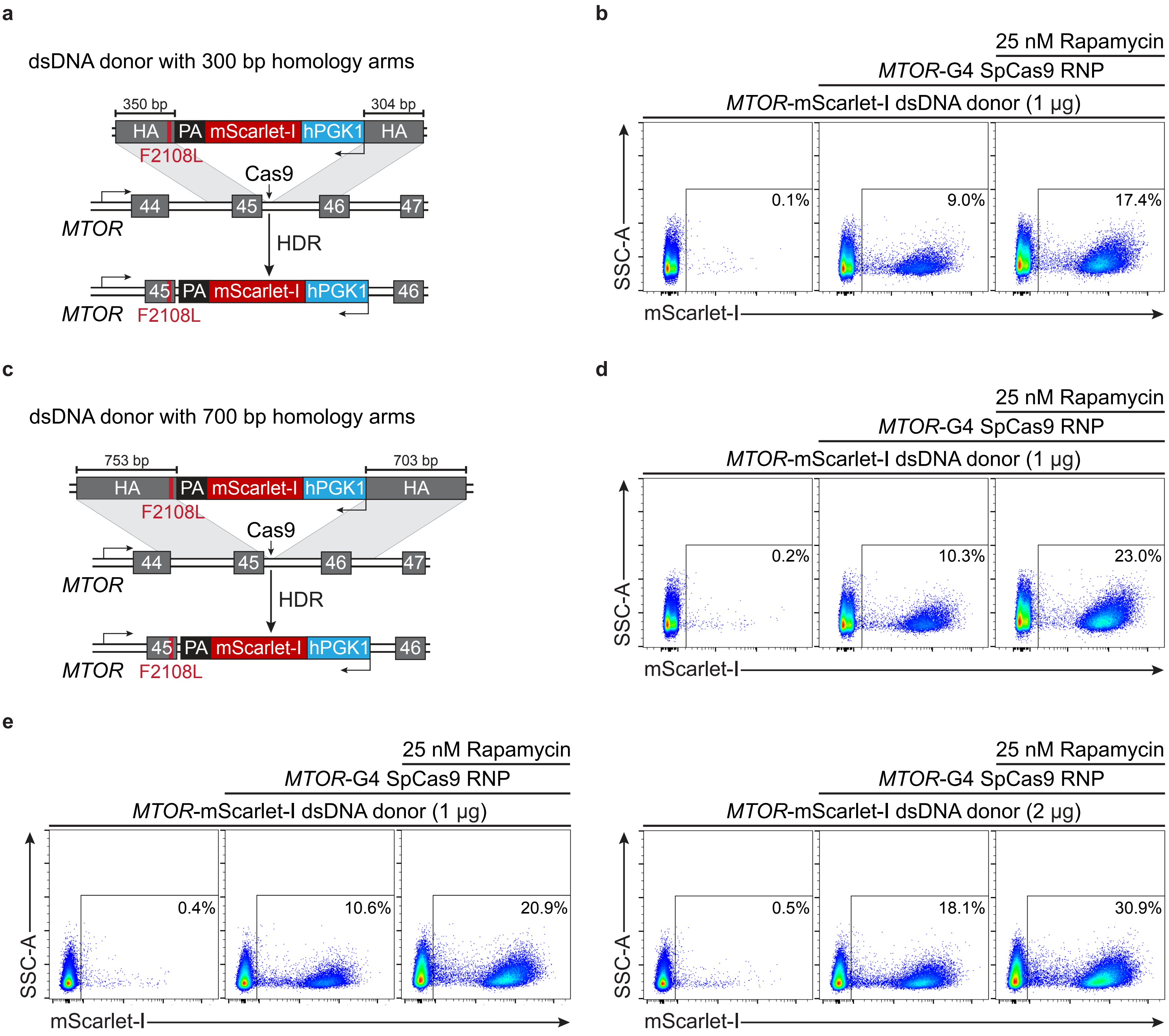


**Figure S5**
**Levesque et al.**

**Supplementary Figure 5. Targeting a fluorescent reporter to the *MTOR* locus allows the selection of rapamycin-resistant $CD3^+$ T cells.** (**a**) Schematic representation of mScarlet-I targeting to the reverse DNA strand of the *MTOR* locus using a linear dsDNA donor harboring 304-350 bp homology arms. The F2108L mutation is introduced via the left homology arm (HA) and transgene expression is driven by a human *PGK1* promoter. (**b**) FACS-based quantification of targeted mScarlet-I integration. $CD3^+$ T cells were transfected with SpCas9-G4 RNP and the indicated amount of linear dsDNA donor and treated with 25 nM rapamycin for 15 days starting 3 days post-transfection. $n = 1$ experiment. (**c**) Same as in (**a**) with a linear dsDNA donor harboring 703-753 bp homology arms. (**d**) Same as in (**b**) with the linear dsDNA donor harboring 703-753 bp homology arms. (**e**) Same as in (**b**) using two doses of the linear dsDNA donor harboring 703-753 bp homology arms and $CD3^+$ T cells treated for 8 days starting 3 days post-transfection. $n = 1$ experiment. *hPGK1*, human phosphoglycerate kinase 1 promoter. PA, polyadenylation signal. HA, homology arm.

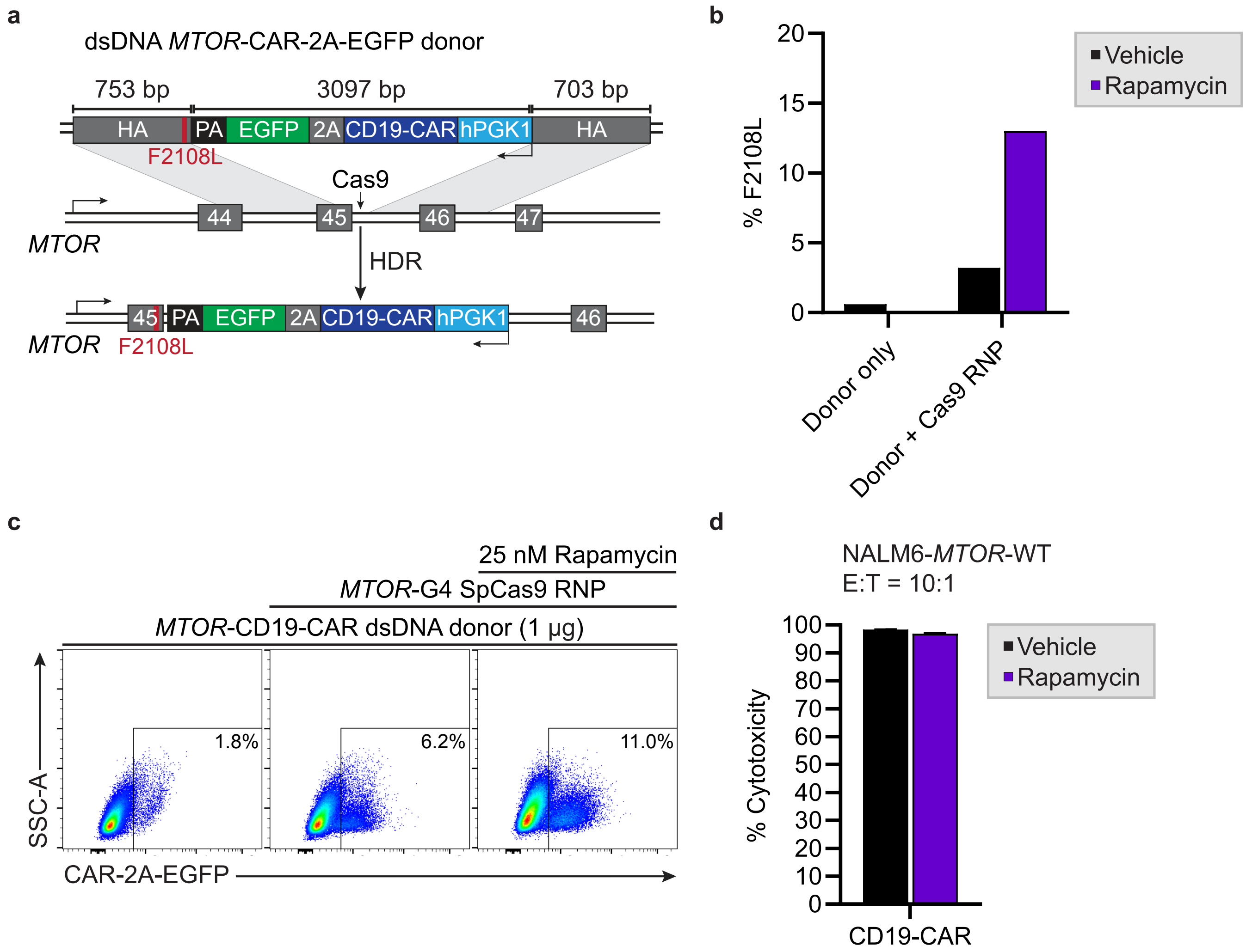


**Figure S6**
**Levesque et al.**

**Supplementary Figure 6. Non-viral targeting of a CAR to the *MTOR* locus allows the selection of rapamycin-resistant CAR-T cells.** (**a**) Schematic representation of CD19-CAR-2A-EGFP targeting to the reverse DNA strand of the *MTOR* locus using a linear dsDNA donor. The F2108L mutation is introduced via the left homology arm (HA) and transgene expression is driven by a human *PGK1* promoter. (**b**) Quantification of *MTOR*-F2108L alleles as determined by TIDER analysis from Sanger sequencing. T cells were treated (rapamycin) or not (vehicle) with 25 nM rapamycin 3 days post-transfection for 8 days. $n$ = 1 experiment. (**c**) Same as in (**b**), but FACS-based quantification of targeted CD19-CAR-2A-EGFP integration. (**d**) Luciferase-based cytotoxicity assay. Following rapamycin selection, CD19-CAR-T cells were incubated with NALM6 cells stably expressing firefly luciferase (FLUC) and 25 nM rapamycin at an effector to target (E:T) ratio of 10:1. Luminescence was measured after 18 hours of incubation. $n$ = 1 experiment. *hPGK1*, human phosphoglycerate kinase 1 promoter. PA, polyadenylation signal. HA, homology arm.

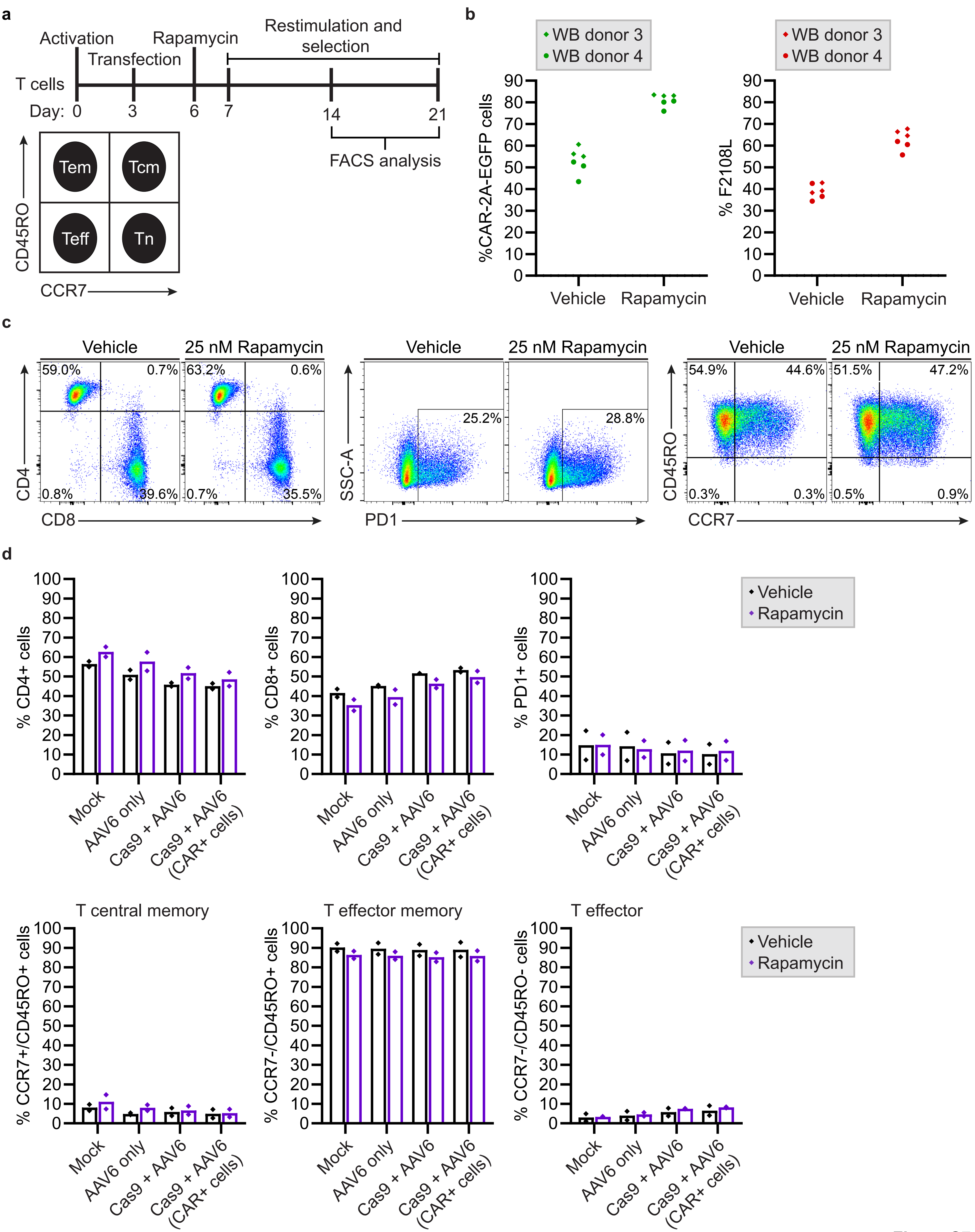


Figure S7
Levesque et al.

**Supplementary Figure 7 (Related to Fig. 3). Rapamycin has minimal impact on T cells' differentiation. (a)** Schematic of the expansion timeline, and representative plot for memory markers. Tn, naïve T cells. Tcm, central memory T cells. Tem, effector memory T cells. Teff, effector T cells. Differentiation and exhaustion markers are analyzed by flow cytometry after 8 days (D14) or 15 days (D21) of treatment with rapamycin. (**b**) CD19-CAR-2A-EGFP knock-in quantification as determined by FACS and TIDER from Sanger sequencing. Whole blood (WB) primary $CD3^+$ T cells were transfected with a SpCas9-G4 RNP and transduced with an AAV6 vector with a MOI of $5x10^3$. T cells were treated (rapamycin) or not (vehicle) with 25 nM rapamycin 3 days post-transfection for 15 days. *n* = 2 independent biological replicates performed in triplicate at different times with $CD3^+$ T cells from two different healthy donors. (**c**) Differentiation and exhaustion markers analysis of the viable CAR-2A-EGFP+ cells after 8 days of treatment with 25 nM rapamycin (D14) as determined by CD4, CD8, PD1, CCR7, and CD45RO staining. (**d**) Same as in (**c**) after 15 days of treatment with 25 nM rapamycin (D21). *n* = 2 independent biological replicates performed in triplicate at different times with $CD3^+$ T cells from two different healthy donors.

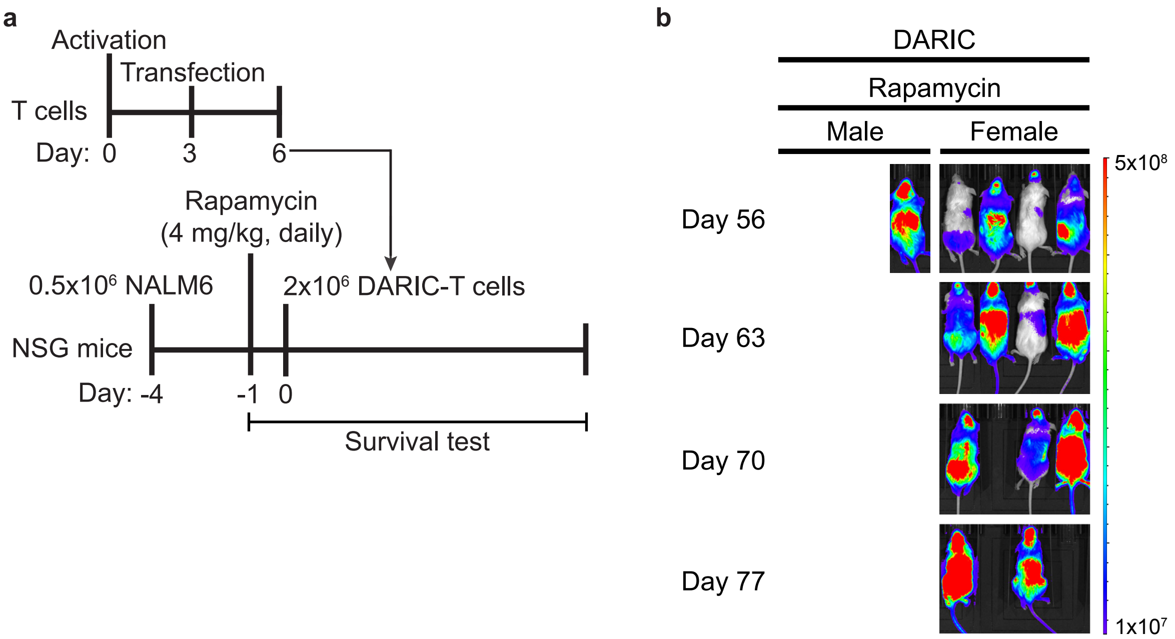


**Figure S8**
**Levesque et al.**

**Supplementary figure 8. Prolonged tumor control in female mice treated with rapamycin and DARIC-T cells (Related to Fig. 6).** (**a**) Timeline and experimental setup for the pre-B acute lymphoblastic leukemia xenograft mouse model. Male and female NSG mice were challenged with 0.5 x $10^5$ NALM6-RFP-FLUC cells and daily rapamycin treatment (4 mg/kg) started three days later. Four days after tumor inoculation, 2 x $10^6$ untransduced (UT) T cells or DARIC-T cells were injected. (**b**) Bioluminescence (BLI) and tumor bio-distribution of mice treated with rapamycin and DARIC-T cells over eleven weeks (See **Fig. 6c** for BLI quantification and tumor bio-distribution of all mice over the first seven weeks). The color barcode represents the radiance scale (photons/s/$cm^2$/sr). $n$ = 8 mice per group.
